# Identifying and quantifying the contribution of maize plant traits to nitrogen uptake and use through plant modelling

**DOI:** 10.1101/2024.04.18.589717

**Authors:** Jie Lu, Tjeerd Jan Stomph, Guohua Mi, Lixing Yuan, Jochem Evers

## Abstract

Breeding for high nitrogen use efficient crops can contribute to maintaining or even increasing yield with less nitrogen. Nitrogen use is co-determined by N uptake and physiological use efficiency (PE, biomass per unit of N taken up), to which soil processes as well as plant architectural, physiological and developmental traits contribute. The relative contribution of these crop traits to N use is not well known but relevant to identify breeding targets in important crop species like maize. To quantify the contribution of component plant traits to maize N uptake and use, we used a functional-structural plant model. We evaluated the effect of varying both shoot and root traits on crop N uptake across a range of nitrogen levels. Root architectural traits were found to play a more important role in root N uptake than physiological traits. Phyllochron determined the structure of the shoot through changes in source: sink ratio over time which, in interaction with light and temperature, resulted in a significant effect on PE and N uptake. Photosynthesis traits were more relevant to biomass accumulation rather than yield, especially under high nitrogen conditions. The traits identified in this study are potential targets in maize breeding for improved crop N uptake and use.

**Highlight:** Our research provides insight into the relevance of a range of traits for maize N uptake and N use, and identifies several potential target traits based on underlying mechanisms to assist maize breeding.

## Introduction

In the past five decades, nitrogen fertilizer application has increased globally along with an increased requirement for food production (Glass, 2003; De Vries et al., 2013). However, more than 60 % of the nitrogen applied is lost through leaching, surface run-off, denitrification and volatilization (Raun and Johnson, 1999; Wang and Li, 2019). For maize, like for many other crops, the maximum production per unit of N fertilizer applied is considered to be genotype-specific (Tsai et al., 1992). Therefore, breeding for high overall crop nitrogen use efficiency at reduced nitrogen inputs is one of the approaches to allow farmers to reduce nitrogen fertilizer input while maintaining high production levels.

Overall crop nitrogen use efficiency is a complex trait and the combined result of several underlying processes during crop growth. Nitrogen use efficiency can be decomposed into two main components: nitrogen uptake efficiency and a measure for utilization efficiency of nitrogen called physiological efficiency (PE) (Van Keulen, 1982). Nitrogen uptake efficiency is expressed as kg N taken up per kg N in the soil; PE is the efficiency with which the N taken up is converted into grain biomass, in kg grain per kg N taken up. In maize, major genetic variation in nitrogen use efficiency under non-limiting nitrogen conditions relates to differences in nitrogen uptake efficiency. Under limiting nitrogen, the genetic variation in nitrogen use efficiency is mainly determined by differences in PE (Gallais and Coque, 2005; Xu et al., 2012; Li et al., 2015). However, it is still unclear which plants traits are underlying the observed difference in PE and N uptake efficiency.

A range of root system architectural, anatomical and physiological traits are associated with nitrogen uptake (Mi et al., 2010, 2016). Fewer crown roots increased 95% of rooting depth located from 26.7cm to 34.4cm in a field experiment in South Africa. This increase of rooting depth contributed to more nitrogen uptake from deeper soil layers and was found especially beneficial under low nitrogen conditions (Saengwilai et al., 2014b). A lower number of crown roots has been associated with a larger nodal root diameter. Fewer but thicker roots were found to result in better growth in a low N environment (Schneider et al., 2021). In addition, increasing the amount of root cortical aerenchyma (RCA), which is related to root tissue density (Postma and Lynch, 2011; Saengwilai et al., 2014a) can increase the rooting depth by reducing metabolic costs of roots through reducing respiration. Besides architectural and anatomical traits, physiological traits such as those related to both high and low affinity transporters of nitrate play a role in nitrogen uptake (Lazof et al., 1992; Parker and Newstead, 2014; York et al., 2016). In experiments, those architectural and physiological traits usually interact, depending on the genotype that is used, and display plastic responses to environmental factors. Therefore, it is difficult to quantify experimentally the effects of individual traits on efficient N use through their effect on N uptake efficiency.

Traits related to shoot architecture, leaf photosynthetic traits and nitrogen reallocation further contribute to nitrogen use efficiency by affecting PE. Traits such as leaf angle distribution along the stem and leaf number determine light distribution within the canopy and as a consequence light interception, leaf area, plant height and yield accumulation (Huang et al., 2017). Photosynthetic traits such as the maximum photosynthesis rate of leaves also differ among maize cultivars (Chen et al., 2013). Maximum photosynthesis rate can be linked with leaf nitrogen content since leaf nitrogen is the major component to construct photosynthetic enzymes such as rubisco (Vos et al., 2005; Mu et al., 2016). When grains start growing, there is competition for nitrogen between grains to form the protein for the next generation and leaves to continue photosynthesis. The ability to maintain a high leaf photosynthesis rate by keeping rubisco levels high is called “stay green”. “Stay green” cultivars tend to have a higher yield combined with a reduced nitrogen remobilization efficiency and a reduced protein concentration in the seeds (Masclaux-Daubresse et al., 2010; Mi et al., 2003). These traits determine PE, and experimentally it is hard to disentangle their effects on both components of PE and maize crop performance.

Crop growth models can help to reveal to which extent individual plant traits contribute to nitrogen use efficiency (Semenov et al., 2007; Asseng et al., 2001). However, in most traditional crop models many plant traits and processes relevant to PE are not explicitly included. Here a modelling approach is adopted, called functional-structural plant (FSP) modelling, that does allow to include such traits and related processes. FSP models simulate stand performance based on individual plants, their growth, physiological functioning and above and below ground 3D architecture when growing in a crop stand and competing with each other for resources (Vos et al., 2010; De Vries et al., 2021). The objective of this study was to quantify the contribution to N uptake and PE of architectural and physiological traits that vary among maize genotypes and in relation to soil nitrogen conditions, using the FSP modelling approach.

## Methods

We expanded an existing whole plant FSP model (De Vries et al., 2021; Evers and Bastiaans, 2016) by including plant and soil processes related to nitrogen uptake and utilization. We then used experimental data from six maize cultivars released in different years between the 1970s and the 2000s grown in a field in 2011 to parameterize the model for maize and the experimental data from these six maize cultivars grown in the field in 2010 to evaluate the model. After that, we performed a series of simulation analyses to quantify the contribution of a range of plant traits to N uptake and PE, especially the traits with great potential for further study.

### Model description

Our 3D maize FSP model represents development and growth of both root and shoot, as a function of competition for light and soil nitrogen within and between plants (Fig. 1; Fig. S1; Fig.S2). Since our goal was to analyze plant behavior, rather than to predict crop performance for specific weather and/or soil conditions like in crop models, in our model the representation of daily light and temperature conditions and the dynamics of soil nitrogen is simpler than in such typical crop models (details below). The model was developed in the GroIMP platform (Hemmerling et al., 2008). To represent maize root systems, seminal roots are initiated as a function of thermal time and the diameters of different root orders are treated as fixed values based on Pagès et al. (2014). Second order (lateral roots on axial roots) and third order (axial roots) root segments are explicitly simulated in 3D while the higher order roots (i.e., lateral roots on lateral roots) are only numerically simulated as carbon sinks but is not considered to enhance N uptake (De Vries et al., 2021), due to computational demand of including higher order roots explicitly. Thus, the model simulates mechanistically plant shoot and root development and growth as driven by respectively temperature, light and soil N and plant carbon and nitrogen sink-source relationships in 3D.

**Figure 1:**
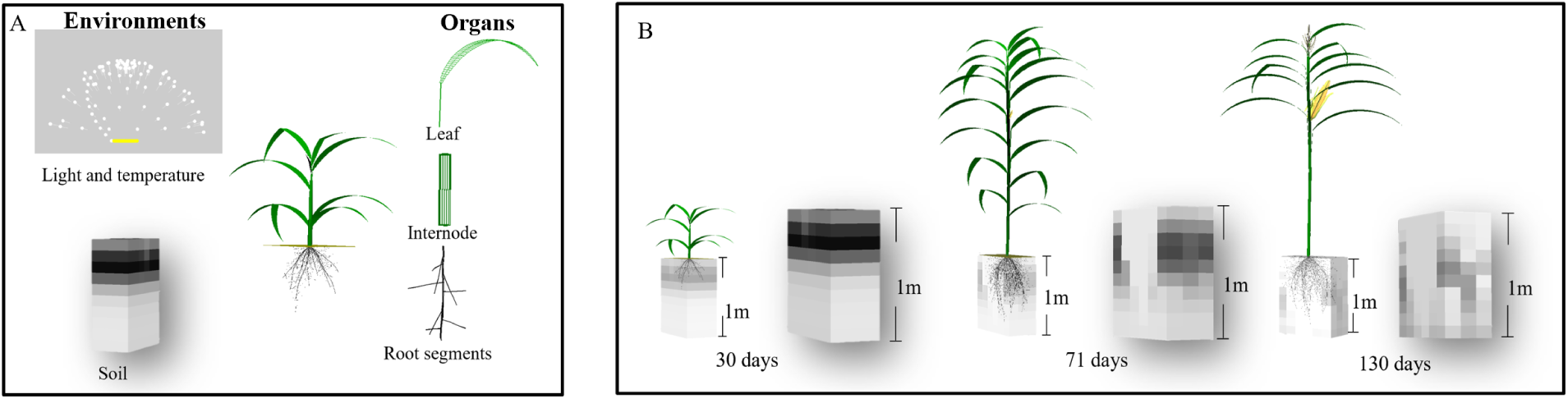
Overview of the maize FSP model including the environmental components (soil cells, temperature and light) and organ types (individual leaf, internode and root segments) of the plant model (A) and model output in terms of plant growth and remaining N in the soil profile at three moments during the maize plant development (B). In the soil cells, darker soil cell colors represent cells with higher soil N concentration. Days are days after emergence.

Light response curves were adopted here to simulate photosynthesis at the leaf level as a function of light absorbed by the leaf. After assimilation organic carbon is allocated to each organ to simulate organ growth based on individual organ sink strength which is defined as the sigmoid function of organ potential dry mass and growth duration (Yin et al., 2003). Therefore, unlike in crop models, the leaf expansion rates in our model are co-determined by leaf sink strength, available carbon and thermal time, in a same manner stem, root and grain growth are modeled. Grain growth starts after the initiation of the final leaf. Root carbon sink strength is considered a fraction of leaf sink strength (De Vries et al., 2021). Non-structural root carbon is considered to be negligible ( 1% of root dry weight) in our model (Wu et al., 2019). Nitrogen, taken up from the soil and reallocated from senescing leaves, is an important component for photosynthetic enzymes. Therefore photosynthetic nitrogen is a crucial determinant of leaf photosynthetic capacity and linked to the light response curve in our model. The N remaining after the photosynthetic nitrogen and the amount of nitrogen required as structural N for organ growth is allocated is stored in a N pool (*Nsource*, g*/*plant). This nitrogen stored in the N pool is available in the next time step for allocation to leaves, stems and grains.

#### Light and temperature in model

Actual daily light and temperature data were obtained from the Jilin weather station. Since the daily data with erratic temperature and radiation values could easily obscure the subtle effects we aimed to quantify (Li et al., 2021), we used as input the annual course in average daily temperature (*T_avg_*, eq. 1) fitted to the measured daily average temperatures at the experimental site in 2010. *Dy* represents the day of the year.

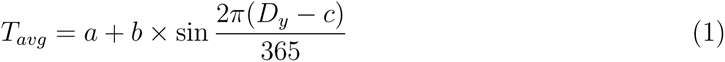

Where *a* (°C) is the yearly average temperature, *b* (°C) is the temperature amplitude and *c* is the day of year with the daily average temperature.

In a similar way the annual course in incoming daily radiation was used as a function of the latitude and day of the year and transmissivity of the atmosphere (Spitters, 1986; Evers et al., 2010), fitted to the daily incoming radiation data from 2010.

To simulate plant growth under field conditions, direct incoming radiation was simulated by using an array of 24 directional light sources based on latitude and day of the year while diffuse light was simulated by using an array of 72 directional light sources positioned in a hemisphere (Evers et al., 2010; Buck-Sorlin et al., 2011). Light capture was simulated by the reverse Monte-Carlo ray tracing algorithm in GroIMP (Hemmerling et al., 2008).

#### Thermal time and plant development

The phenological development of maize plants from sowing to maturity was expressed as a function of thermal time (*tt*) (°Cd). Thermal time (eq. 2) in the model was calculated from daily average temperatures (*T_avg_*, °C) and a maize base temperature (*T_b_*, °C). *T_b_* was set to 10 °Cd for these maize cultivars (Chen et al., 2013). No optimum temperatures for development where considered as in the experiments used to calibrate and check the model daily maximum temperatures rarely were above reported optimum temperatures for development of maize (30.8 °Cd) (Sáanchez et al., 2014).

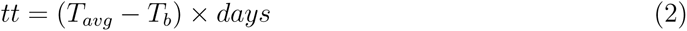

#### Soil N

The soil was composed of cubic soil cells of which the size (*Cellsize*, m) was 0.1x0.1x0.1 m. Simulated soil depth was set to 1 m to account for N percolating beyond the rooting zone as observed rooting of the simulated cultivars was maximally 0.6 m. No further root biomass accumulation was observed after silking in the field experiment (Shao et al., 2018). Therefore, root elongation was assumed to stop when all leaves were fully expanded. In contrast to many maize cultivars, root segments are rarely found beyond 0.6 m depth for the maize cultivars and growing conditions for which we conducted the analyses (Ning et al., 2014). As all data used for verification was under no-water-stress conditions and for simplicity, we did not explicitly consider the effects of precipitation or soil temperature.

For individual soil cells at each simulation step (*t*, day), soil nitrogen (*SoilN* (*t*), *µ*mol*/*m^3^) was a function of the amount of N applied at the start of the simulation (*Napp*, *µ*mol*/*m^3^), the additional N application (*Napp*2, *µ*mol*/*m^3^) at certain date (*DNapp*2), the initial soil N (*Ninit*, *µ*mol*/*m^3^), mineralization (*Nm*, *µ*mol*/*m^3^) and leaching (*Nl*, *µ*mol*/*m*^−^*^3^) and uptake by the roots present in the cell (*Nu*,*µ*mol*/*m^3^). The *SoilN* (*t*) is described in eq. 3 (Fig. S3).

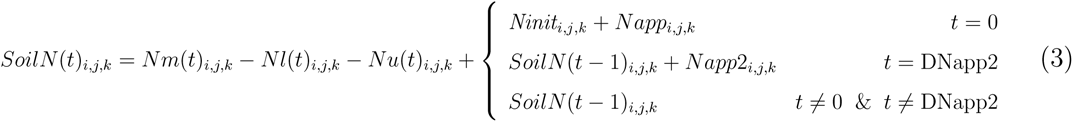

Where *i*, *j*, and *k* are the Cartesian coordinates of a soil cell, of which *k* indicates the vertical coordinate, and *Cvol* (m^3^) is the cell volume. The initial and applied N were assumed to be distributed homogeneously in the top 0.3 m soil. The initial amount of N in the soil cells deeper than 0.3 m were set to 0.

The net N change at any point in time in a soil cell at depth *k* through leaching (*Nl*(*t*)*_i,j,k_*) was always the sum of loss to the cell at depth *k* + *Cellsize* directly below, and gain from leaching from the soil cell at depth *k − Cellsize* directly above. To reduce the computing time, no lateral N movement between soil cells was considered. Leaching finally accumulated in the bottom soil layer from which no leaching loss was calculated (eq 4, 5).

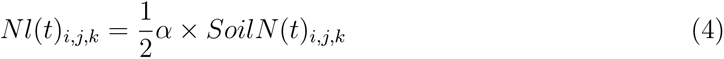

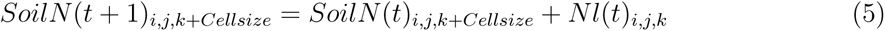

where α is soil permeability to water and hence to nitrogen (Addiscott and Whitmore, 1991).

To simulate N made available by mineralization in the top 0.3 m soil (Stanford and Smith, 1972), organic nitrogen (Norganic, *µ*mol*/*m^3^) was assumed to gradually mineralize and provide extra N that can be taken up by the root (eq. 6, 7, 8).

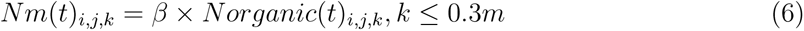

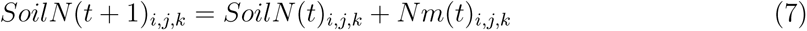

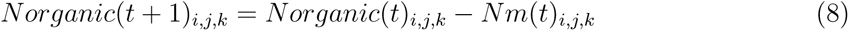

where *β* is the relative mineralization rate of available soil organic nitrogen.

#### N uptake

N uptake by the root system was calculated as the cumulative uptake across all soil cells (eq. 13). For each soil cell separately N uptake was calculated for the root segments in that soil cell. For per root segment uptake, Michaelis-Menten kinetics were used to simulate N uptake through the high affinity transport system (*HATS*, *µ*mol*/*(m^2^ *·* day), Eq. 9) and a linear relation with soil N to simulate N uptake through the low affinity transport system (*LATS*, *µ*mol*/*(g *·* day), eq. 10) (Siddiqi et al., 1990). eq.12 represents nitrogen uptake (*µ*mol*/*day) for each root segment per simulation through the two N transport systems. *E_N_* (dimensionless, eq. 11) represents a negative feedback of plant nitrogen concentration on root nitrogen uptake by the combined transport systems. Based on previous studies, this negative feedback was added to N uptake in the model (Forde and Clarkson, 1999; Bertheloot et al., 2011; Barillot et al., 2016).

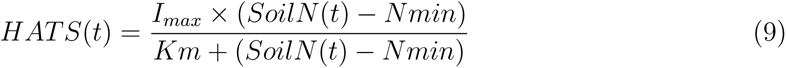

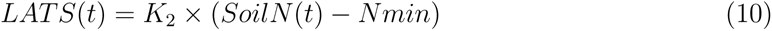

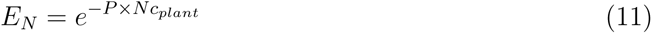

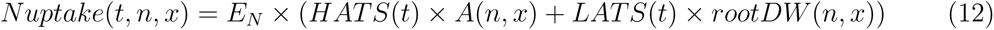

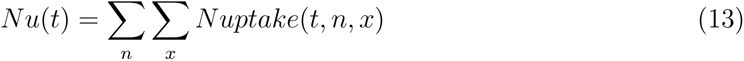

Where x is the root segment rank within a root and n is the root primordium number. *Nmin* (*µ*mol*/*L) is the minimum N concentration for nitrogen uptake. *I_max_* ( *µ*mol*/*(m^2^ *·* day)) is the maximum influx rate of N. *Km* ( *µ*mol*/*m^3^) is the soil nitrogen concentration at half *I_max_*. Both parameter values were derived from York et al. (2016). *K*_2_ (*µ*mol*/*(g *·* day)) is the constant rate of *LATS* activity, which was derived from Pace and McClure (1986). *A* (m^2^) is the surface area of an individual root segment and *rootDW* (g) is the dry weight of the individual root segment. *P* (dimensionless) is the coefficient for the negative feedback of plant N concentration on root nitrogen uptake. *Nc_plant_* (g N*/*g DW) is the whole plant nitrogen concentration.

A certain amount of nitrogen is fixed in the root to form root structure to support growth of the root system. This amount of nitrogen is assumed not-available for the shoot. In the model, it is a fixed fraction (*f Nroot*, g N*/*g DW) of the root dry weight of the plant accumulated during each simulation step. The rest of the nitrogen is available to the shoot (eq. 15). When during a time step the amount of newly acquired and stored N cannot fulfill the N demand for growth of the root, all N is fixed in the root and none of the N taken up in a time step becomes available to the shoot. This leads to a reduction in the shoot N concentration. In case N was available as stored N from the previous time step this is partly allocated to the root and partly to the shoot based on their relative sink strength for N.

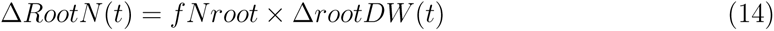

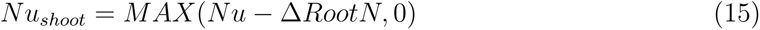

#### Grain nitrogen

Once the plant enters the reproductive stage, part of the available nitrogen is allocated with priority to the grains (Masclaux-Daubresse et al., 2008, 2010), using a linear relationship between leaf N and grain N concentration in maize (Fernandez et al., 2021). Under low nitrogen, the grain nitrogen influx rate was less than the rate under sufficient nitrogen (Ning et al., 2017). Therefore, to account for balancing the plant N flux between photosynthetically active leaf tissue and grain protein accumulation, the fraction of nitrogen (*f Ngrain*, g N*/*g DW) allocated to the grains was considered to be a function of leaf nitrogen concentration from the previous time step (*Nleaf* (*t −* 1), g N*/*g DW).

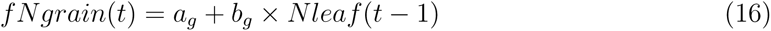

The total N in the grains at any point in time (*GrainN* (*t*)) is the accumulation of

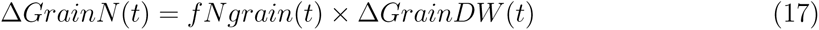

*a_g_* is the intercept of the relation between *f Ngrain* and *Nleaf* , representing at which leaf nitrogen concentration no nitrogen is allocated to the grains. *b_g_* is the slope of this relation.

#### Photosynthetic nitrogen

During the vegetative stage, the nitrogen available to the shoot functions as photosynthetic nitrogen and is distributed over the leaves and internodes (*StemN_photo_*, g) according to their nitrogen demand. The organ (i.e. leaf and internode) demand for photosynthetic nitrogen (*DNarea_photo_*, g*/*m^2^) is the smallest of either 1) the nitrogen demand (*DNareal_photo_*, g*/*m^2^) as a function of the light gradient in the canopy and the target nitrogen concentration (*LN* 0, *g/m*^2^) of fully lit leaves (eq. 18) (Hikosaka et al., 2016) or 2) the nitrogen demand (*DNareaa_photo_*, g*/*m^2^) as a function of the individual organ’s age (*AgeD*, day) (eq. 19).

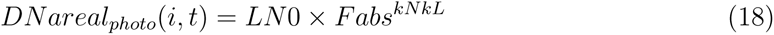

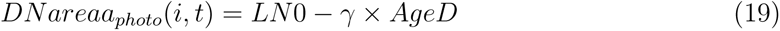

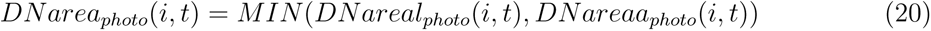

Where i represents an individual photosynthetic organ (i.e. individual leaf or internode). *Fabs* is the fraction of incoming light absorbed by that organ. *kNkL* is the ratio between nitrogen and light extinction coefficients, and set equal to 0.368 (Hikosaka et al., 2016). *γ* is the coefficient of demand of photosynthetic nitrogen controlled by age and set equal to 0.02 (Chen and Mi, 2018).

Similar to the carbon sink and source relationship in the model, we integrated a nitrogen sink and source relationship for photosynthetic nitrogen. If available organ photosynthetic nitrogen (*avaN_photo_*, g) was more than organ demand (*DLN* , g), the amount of nitrogen allocated to each photosynthetic organ was equal to its demand for nitrogen. The surplus nitrogen was then assumed to be stored in the stem as a part of a *Nsource*. If the total amount available was less than the total demand, the amount available was allocated over all organs, based on the organ nitrogen demand times the relative nitrogen sink strength of that organ (eq. 21).

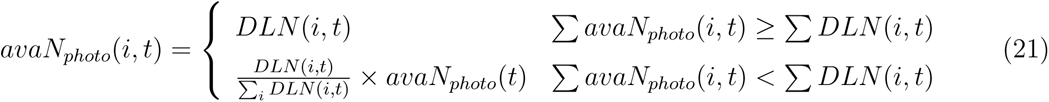

As above, *i* represents an individual photosynthetic organ (leaf and stem) and *t* is the unit time (*day*).

An asymptotic exponential relationship was used to estimate photosynthesis rate (eq. 22). This relationship is widely used in different crop models, such as e.g. SUCROS (Archontoulis and Miguez, 2015):

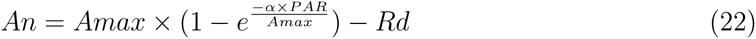

where *Amax* (*µmol CO*_2_*/*(*m*^2^*·s*)) represents the maximum photosynthetic rate, *α* represents the initial slope of the curve at low irradiance levels, *Rd* (*µmol CO*_2_*/*(*m*^2^ *· s*)) is the dark respiration rate and *PAR* (*µmol/*(*m*^2^ *· s*)) represents photosynthetically active radiation.

The *Amax* of photosynthetic active organs was related to specific leaf (or stem) nitrogen (*SLN* , g*/*m^2^) through a modified logistic model (Sinclair, 1989; Vos et al., 2005).

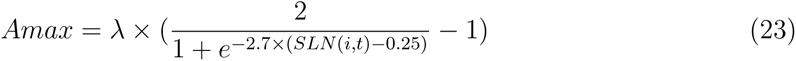

Where *λ* (*µmol/*(*m*^2^*·s*)) is the maximum value of *Amax* under non-limiting *SLN* . *SLN* (*i, t*) is calculated by *avaN_photo_*(*i, t*) and the surface area of the *i*th leaf or internode.

#### Stem nitrogen

During the stem extension period, a proportion of the available nitrogen is allocated to the internodes to maintain a fixed internode N mass fraction (*f Nstem*, g N*/*g DW). This nitrogen is considered to be incorporated in the stem structure and therefore cannot be reallocated. Additionally, stems contain nitrogen for photosynthesis (*StemN_photo_*, see section Photosynthetic nitrogen) which is assumed to be available for reallocation when the generative phase has started (Ning et al., 2017). Since the stem is also assumed to act as a buffer organ, N stored in the pool for later use (*Nsource*, g*/*plant) is also assumed to be contained in the stem until required elsewhere:

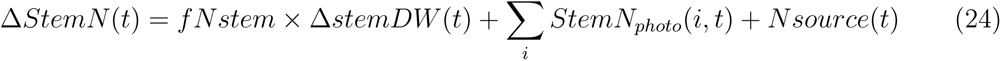

#### Leaf nitrogen

All nitrogen in leaves is considered to contribute to photosynthesis, including leaf structural nitrogen. When leaf nitrogen drops below a critical leaf nitrogen (*Leaf N_critical_*, g*/*m^2^), the leaf sheds regardless of whether it reached its life span. Once a leaf is shed, all nitrogen except for leaf structural nitrogen (*Leaf N_struct_*, g*/*m^2^) is available for use elsewhere and photosynthesis of that leaf stops. Both values of leaf structural nitrogen and leaf critical nitrogen were derived from Chen et al. (2016).

#### Carbon allocation between root and shoot

Based on the functional equilibrium theory (Lambers, 1983; Shipley and Meziane, 2002), plants systematically invest extra carbon when exposed to resource limitation, to support the growth of organs relevant to acquire the more limiting resource more effectively to optimize whole-plant growth. This concept is adopted here by implementing a relationship between root-to-leaf biomass partitioning ratio (*rlratio*) and *Nleaf* , which was assumed constant across plant developmental stages. This relationship was implemented in the model to control carbon allocation between shoot and roots for each time step (See Model description section) in response to leaf N levels.

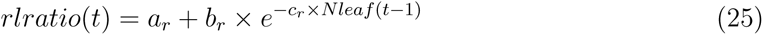

where *a_r_*represents the minimum value of *rlratio*, and *b_r_* and *c_r_* are the coefficients for the negative feedback between leaf N concentration and *rlratio* (Fig. S4, Fig. S5). The coefficients were derived for the two cultivars (Zhengdan958 and Xianyu335) on which we had most information and are for this study considered cultivar independent as no clear difference were found between these two cultivars.

### Field experiment

A field experiment was conducted in Siping (43°17’N, 124°26’E), Jilin province, China, in 2010 and 2011, reported in Chen et al. (2013, 2014). A split plot design was used with 4 replications in 2010 and 5 replications in 2011. Nitrogen treatments were the main plot and genotypes were the subplots. The selection of maize cultivars used to parameterize and evaluate the model were Zhengdan958 (ZD958), Xianyu335 (XY335), Zhongdan2 (ZD2), Danyu13 (DY13), Yedan13 (YD13) and Nongda108 (ND108) which were all released between the 1970s and the 2000s in China. Three nitrogen levels were applied: 240kg/ha (HN), 120kg/ha (MN) and 0kg/ha (LN). For HN and MN, half the nitrogen was applied before sowing and the other half was applied at the V8 stage (the eighth leaf fully expanded). Maize seeds were hand-sown at a density of 6 plants*/*m^2^ and a row and plant distance of 0.6 and 0.28 m, respectively. The crops were rain-fed. Precipitation was 580 and 401 mm in 2010 and 2011, respectively (Chen, 2015). Biomass and nitrogen concentration for each organ (leaf, stem and grain) were collected separately for three plants of each cultivar within each nitrogen treatment and replicate, at both silking and maturity stages in both years. Biomass was oven dried at 70 °C until dry weight was constant. N concentration of separate organs (i.e. leaf, stem and grain) was then measured by the semi-micro Kjeldahl method (Nelson and Sommers, 1973).

Photosynthesis light response curves were derived for ear leaves of the six maize cultivars per plot at grain filling stage under HN using a portable photosynthesis system (Li6400; LI-COR, Lincoln, NE, USA). The light intensity was set to 0, 20, 50, 100, 200, 500, 1000, 1300, 1600 or 2000 *µmol/*(*m*^2^ *· s*). After measuring photosynthesis rates, the ear leaf was removed, dried and measured for biomass, leaf area and nitrogen mass concentration.

At grain filling stage, roots of the six cultivars grown at HN and LN in 2011 and at HN in 2010 were excavated by sampling a 60 cm wide by 28 cm long by 60 cm deep soil block that was divided into three 20cm layers. Per layer, roots were washed with a 0.4mm sieve and axial roots and lateral roots were separately scanned (Epson1600, India). WinRhizo (version Pro 5.0, Canada) was used to analyze root traits (such as root diameters, root length). After scanning, root samples were dried and weighed.

### Model parameterization

The values of the model parameters were derived from the 2011 experimental data. The cultivar specific parameters *a_g_* and *b_g_* in eq. 16 to define grain nitrogen concentration in response to leaf N concentration were derived from relations fitted by linear regression using the“lm” function in R. Maize grain N concentration is quite stable during grain development (Ning et al., 2017; Fernandez et al., 2021). The data of grain N concentration and leaf N concentration at physiological maturity from the 2011 field experiment were used here to represent the N balance between grain and leaf during grain development.

Nested models (mle2) in the R package “bbmle” were used to derive photosynthesis related parameters. Multiple models were fitted to obtain the best parameter set. The models either assumed that all three parameter values for photosynthesis (eq. 22) differed between cultivars, only two of them differed and one was shared, only one of them differed and two were shared, or all three were shared among cultivars. The Akaike information criterion (AIC) was used to select the best model. By definition, smaller AIC values represent better overall fits with adjustment on parameter number. If the difference in AICs between two models is less than 2, the two models can be considered equivalent and therefore, in this case, the simpler model was chosen (Bolker, 2008). The parameters final leaf number (*Leaf Num*), *seedMass*, *f Nstem*, *initD* were based on direct measured data. Root tissue density (*RTD*) was calculated by dividing root biomass by root length and the average ratio of the diameter of the daughter root to that of the mother root (*RDM* ) was calculated by dividing the average diameter of first order lateral roots by that of the axial roots. Means and standard errors were calculated for each cultivar and listed in Table S2.

### Model evaluation

Both actual daily weather and fitted relations were used to evaluate performance of our model. Temperature and incoming radiation (Table S1) were derived from local weather data during the 2010 growing season. The “nls” function in R version 4.0.2 (Milliken, 1990; R Core Team, 2021) was used here to fit the temperature and incoming daily global radiation patterns for 2010 (eq. 1) to test modelled N uptake, yield and physiological efficiency (PE) versus observed data. PE in this study is defined in eq.26. Simulated root length proportion (rRLD) for every 20cm soil layer were calculated in eq 27.

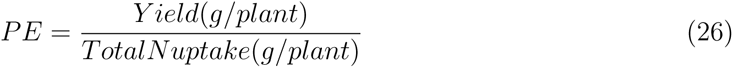

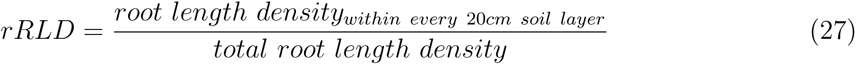

Soil nitrogen leaching and mineralization have been taken into account in this model which means that at each time step the amount of nitrogen in a soil cell is one of the components to determine available soil nitrogen in that cell for a next time step. Therefore, nitrogen fertilizer application as done in the 2010 experiment was adopted in the model (half of the N applied before sowing and half at V8 stage). In addition, based on the recorded dates from 2011, plants were set to emerge 17 days after sowing, for model evaluation purposes. All the environmental and management parameter values are listed in Table S1. Simulations were run under the three different nitrogen treatments (HN, MN and LN) by converting nitrogen applications from the field experiment to model input settings (Table S3). The parameter values of *Ninit* and *Nm* were assumed based on final plant N uptake of unfertilized plants (treatment LN).

Simulation runs were replicated five times to take into consideration the variation in output caused by the light model; an inherently stochastic computation. Each run consisted of a single plant. To reduce border effects, above ground canopy light conditions were created by cloning the plant 10 times in both x and y directions using the replicator functionality of GroIMP. Belowground, periodic boundaries were used so root segments that laterally extended beyond the plot boundary entered the cell on the opposite side (De Vries et al., 2021).

The relative root mean square error (rRMSE) was used to estimate differences between observed and simulated values for N uptake, yield and PE:

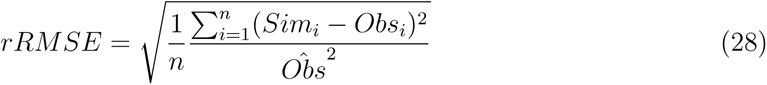

Where *Obs_i_*and *Sim_i_* are the observed and simulated values and n is the number of observed and simulated values.

### Simulation experiment

A simulation experiment was done to quantify the relevance of genotype-specific plant architectural, developmental and physiological traits for N uptake, yield and PE (physiological efficiency). To run the simulation experiment, sowing date was set to Julian day 129 (May 9th) based on the 2010 field experiment and total growth duration was set at 154 days at which grain biomass was stable. The parameter values of temperature and light derived from the 2010 weather data were used as input. Plant density and nitrogen fertilizer application were taken from the 2010 experiment as well. In this simulation experiment, the earlier established parameter values from the cultivar ZD958 were used as default scenario since the parental line of ZD958, Zheng58, is widely used in current Chinese maize breeding program and ZD958 has been well studied in China.

We chose a range of plant traits potentially having a major effect on PE, to test by first individually changing values of root-trait related parameters *initD*, *RDM* , *RootNum*, *RTD*, *I_max_*and *K*_2_ which were all genotype-specific (Table 1) and photosynthesis and leaf growth related parameters: *λ*, *b_g_*, *Leaf Num* (both cases with and without an effect on the time of silking were included), the time between leaf appearance (*phyllochron*) which affects time of silking without affecting leaf number; and finally grain growth related parameters: potential grain biomass (*wmax*, g), grain growth duration (*te*, °Cd), grain growth rate and grain nitrogen influx rate (Table 1). To account for the fact that two plant traits of interest (axial root diameter and time to silking) in the model are the result of component parameters, two combinations of model parameters were used (Table 1). The combination of *initD* and *RDM* represented axial root diameter. The combination of phyllochron and total leaf number represented the final leaf number on the plant with a given time to silking. Not all traits tested appear in the equations presented above, such as LeafNum and Phyllochron. Such parameters directly drive plant development without the need for an equation; this is fundamental to FSP modelling. The complete model code is available online (https://git.wur.nl/lu068/cn-maize) and the full list of parameters for the plant model is listed in Table S5.

**Table 1:** List of plant traits tested in the parameter sensitivity test, their default values and numbers of related equations.

| Description | Related<br>model<br>parameter | Eq. | Unit |
| --- | --- | --- | --- |
| Maximum influx rate of soil nitrogen for HATS activity | Imax | 9 | $\mu\text{mol}/(\text{m}^2 \cdot \text{day})$ |
| Constant rate of LATS activity | $K_2$ | 10 | $\mu\text{mol}/(\text{g} \cdot \text{day})$ |
| Root diameter for whole root system | initD | - | m |
| Axial root diameter | initD and RDM | - | m |
| Lateral root diameter | RDM | - | m |
| Maximum root number | RootNum | - | - |
| Root tissue density | RTD | - | $\text{g}/\text{dm}^3$ |
| Final leaf number | LeafNum | - | - |
| Time lag between two successive leaf appearance | Phyllochron | - | $^{\circ}\text{Cd}$ |
| Silking time | LeafNum and Phyllochron | - | - |
| Maximum photosynthesis under non-limiting leaf nitrogen | $\lambda$ | 23 | - |
| Potential grain dry weight | wmax | - | g |
| Grain growth duration | te | - | $^{\circ}\text{Cd}$ |
| Nitrogen influx rate in response to leaf nitrogen concentration | $b_g$ | 16 | g N/g DW |

### Parameter sensitivity

In order to identify and quantify the effects of changes in any plant trait alone or in combination, we changed the value of the related parameters by +10% and -10% and recorded the change in output parameter values (N uptake, yield, PE). To achieve a +10% change in axial root diameter, we increased the *initD* by +10% and then decreased *RDM* by 10% to keep lateral root diameter the same. To achieve a +10% change in the final leaf number without affecting time to silking, we increased the final leaf number by 10% and then decreased phyllochron by 10% to maintain silking time. The opposite was done to achieve a -10% change in both plant traits.

Sensitivity was calculated as:

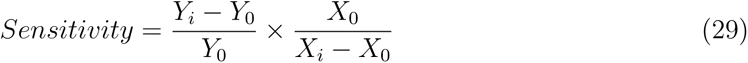

where *Y*_0_ is the output value with default input parameters and *Y_i_* is the output value with a change in input parameters. *X*_0_ represents the default input parameter and *X_i_* represents the input parameter with a change. If the value of sensitivity was larger than 1 or smaller than -1, the output was considered to be sensitive to changes in the input parameter. When values remained between -1 and 1, the output was considered to be insensitive. A positive value meant that when an input parameter increased or decreased by 10%, the change in output values was in the same direction. A negative value meant a change in output in the opposite direction.

Sensitivity was checked for three key output variables: 1) seasonal N uptake (g N/plant), 2) yield (g grain/plant), and 3) PE or physiological efficiency (Grain (g)/N (g)), which is the ratio of yield and N uptake. In addition, root N uptake efficiency was calculated in eq. 30 as a function of changes in Imax and K2.

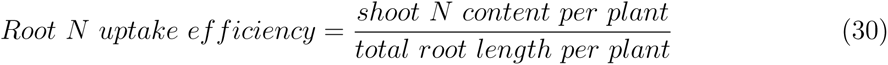

The sensitivity tests were run under high (Napp = 5.714 *µ*mol*/*m^3^) and low (Napp = 0 *µ*mol*/*m^3^) N input conditions to see the potential ability of the tested plant trait values to contribute to N uptake, yield and PE. The plant N concentrations at maturity for both N levels are shown in Fig. S6. The ‘ggplot2’ package (Wickham, 2009) was used to produce figures.

## Results

### Model parameterization

Based on observed daily average temperature and daily global radiation of the growing season in 2010, curves were established as model input (Fig. S7; Table S1). The initial slope of the light response curve at low irradiance levels, *α* (0.0506 *±* 0.0023) and dark respiration Rd (2.43 *±* 0.245 *µ*mol*/*(m^2^ *·* s)) in the photosynthesis model (eq. 22) were the same for all cultivars (Fig. S8; Table S4). In contrast, the *λ* in the relation of *Amax* to specific leaf nitrogen (eq. 23) was found cultivar specific. In the model, *λ* was set from 21.2 for ND108 to 31.8 for XY335 based on field measurement of ear leaf photosynthesis under HN in 2011 (Table S2).

The parameters *a_g_* and *b_g_*, (eq. 16) defining the relation between grain and leaf nitrogen concentration were also found cultivar specific parameters (Fig. S9). On the basis of the 2011 field data (Table S2), *a_g_* ranged from -0.0128 (XY335) to 0.00621 (ND108) and *b_g_* from 0.433 (ND108) to 1.97 (XY335).

### Model evaluation

Using daily temperature and global incoming radiation to evaluate the model gave correspondence between modelled and observed N uptake (rRMSE= 0.190), yield (rRMSE= 0.147) and PE (rRMSE= 0.125) (Fig. S10). With the fitted daily temperature and global incoming radiation, we found a comparable overall correspondence between modelled and observed N uptake with rRMSE equal to 0.233 (Fig. 2A). Interestingly, the model slightly underestimated the N uptake for certain cultivars under low N, while overestimated N uptake for other cultivars under high N. This could indicate the current model is lacking potentially relevant processes such as plastic responses to N level. The overall correspondence of simulated and observed yield values was acceptable with rRMSE equal to 0.174 with similar trend as N uptake (Fig. 2B). Simulated and observed physiological efficiency corresponded well with an overall rRMSE of 0.135 (Fig. 2C). The simulated root length proportion across soil layers showed *∼*75% roots were distributed in the top 20cm of the soil profile (Fig. S11).

**Figure 2:**
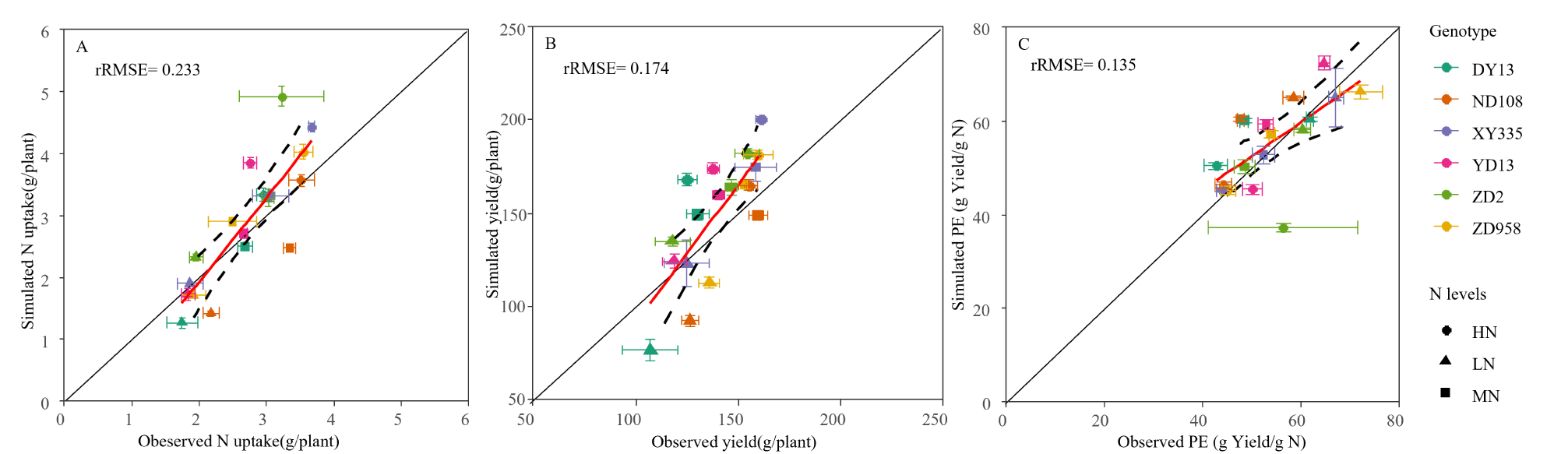
Scatter graphs of simulated versus observed N uptake (A), yield (B) and PE (C) for six maize cultivars grown at high (240kg N/ha, HN), medium (120kg N/ha, MN) and low (0kg N/ha, LN) N fertilizer application rates. Observed data are from experiments in 2010 reported in Chen et al. (2014), where points and error bars represent means *±* SE (n=4). The solid lines represent the fitted line between observed and simulated data while the confidence intervals of the fitted lines were represented by the area between the two broken curves.

### Parameter sensitivity

#### N uptake mechanism

Nitrogen uptake, PE and yield were not sensitive (i.e. sensitivity values were between 1 and -1) to changes in parameter values related to high or low affinity N transport (Fig. 3). Changing the high-affinity related parameter *I_max_* by 10% resulted in sensitivity values ranging from -0.86 to 0.59 for N uptake, yield and PE under both high and low N conditions ( Fig. 3A, B, C). Changing the low-affinity related parameter *K*_2_ by 10% (eq. 10) gave sensitivity values from -0.51 to 0.33 for N uptake, yield and PE under both high and low N conditions (Fig. 3D, E, F). The root N uptake efficiency increased from -1.1% to 2.3% when increasing or decreasing *I_max_*by 10% under both high and low N conditions. The root N uptake efficiency increased from -0.5% to 1.7% when decreasing or increasing *K*_2_ by 10% under both high and low N conditions.

**Figure 3:**
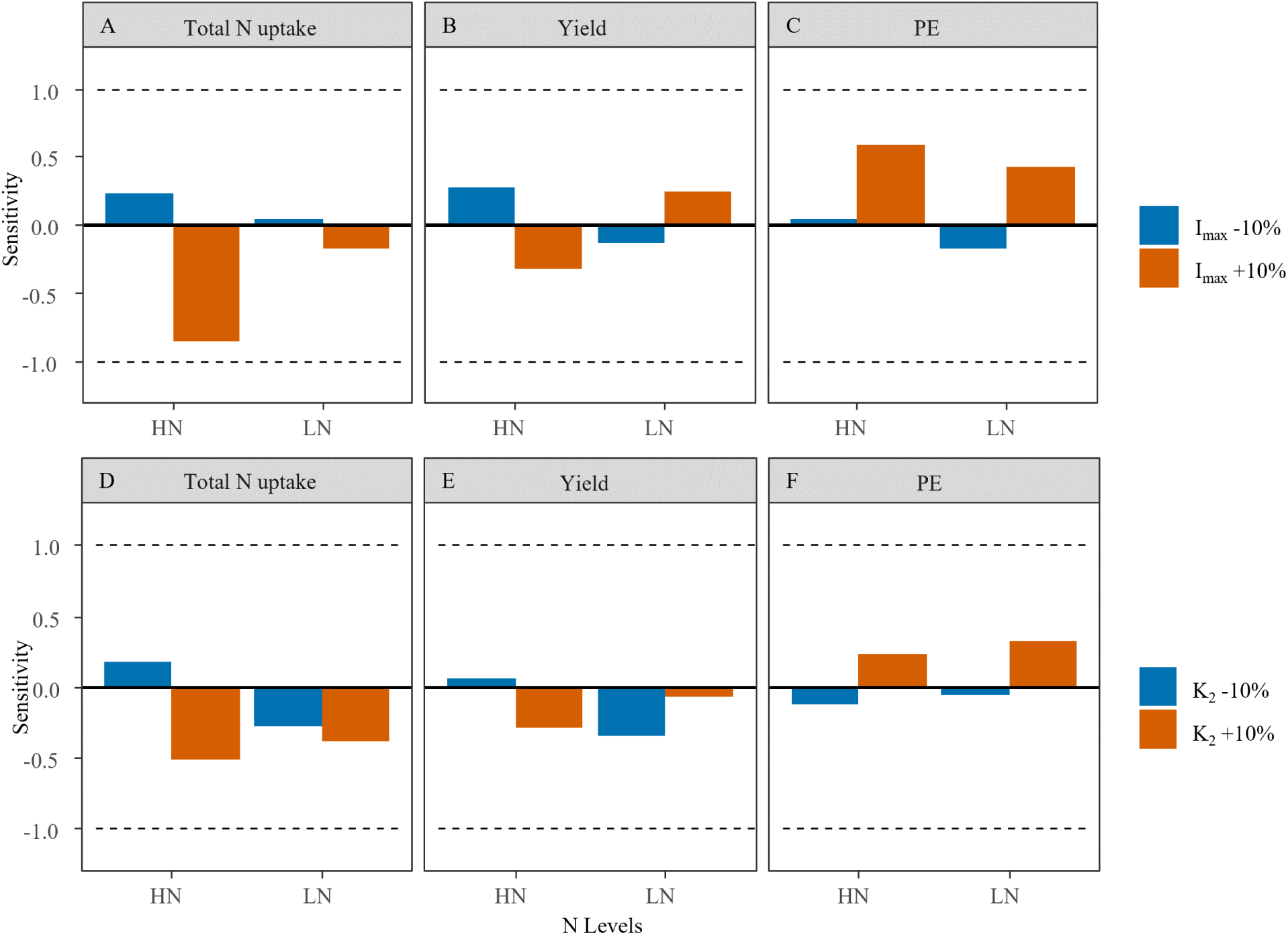
Sensitivity values of total N uptake (g/plant, A, D), yield (g/plant, B, E) and PE (g yield/g N, C, F) under high nitrogen condition (240kg/ha, high N) and low nitrogen condition (0kg/ha, low N) to changes in N uptake capacity related traits: *I_max_*(A, B, C) and *K*_2_ (D, E, F). All analyses were using parameter values of cultivars ZD958 as default. The dashed lines represent model sensitivities of 1 and -1. Parameter and output change in the same direction give a positive sensitivity value; parameter and output change in the opposite direction give a negative sensitivity value. Blue and orange bars represent sensitivity of an output parameter to respectively a 10% decrease or increase in a trait value compared to its default.

#### Root morphology

Nitrogen uptake was sensitive to changes in diameters of all root classes (values between -1.06 and -1.9, Fig. 4A). The negative sensitivity values meant that the response in N uptake was in the opposite direction of the change in parameter value. In other words, reducing root diameter enhanced N uptake more than proportionally, and, increasing root diameter reduced N uptake. When decreasing diameters of all root classes under high N, the change in N uptake was less than proportional (-0.814). For yield a similar major negative sensitivity was observed when changing diameters of all root classes under low N (-1.58 and -1.56, Fig. 4B) meaning that increasing root diameter reduced yield and decreasing root diameter enhanced yield. PE did not show sensitive to changes in diameters of all root classes by 10% under both N levels (from 0.29 to 0.96, Fig. 4C). Nitrogen uptake and yield were more sensitive to changes in diameters of all root classes under low N than under high N (Fig. 4A, B). Only under low N, nitrogen uptake (-1.36) and yield (-1.62) were enhanced by a decrease in axial root diameter alone (Fig. 4D, E). N uptake (-1.02) and yield (-1.50) were sensitive to decreasing only the diameter of secondary and tertiary roots under low N while increasing the diameter of these roots had no major effect on either N uptake or yield (Fig. 4G, H, I).

**Figure 4:**
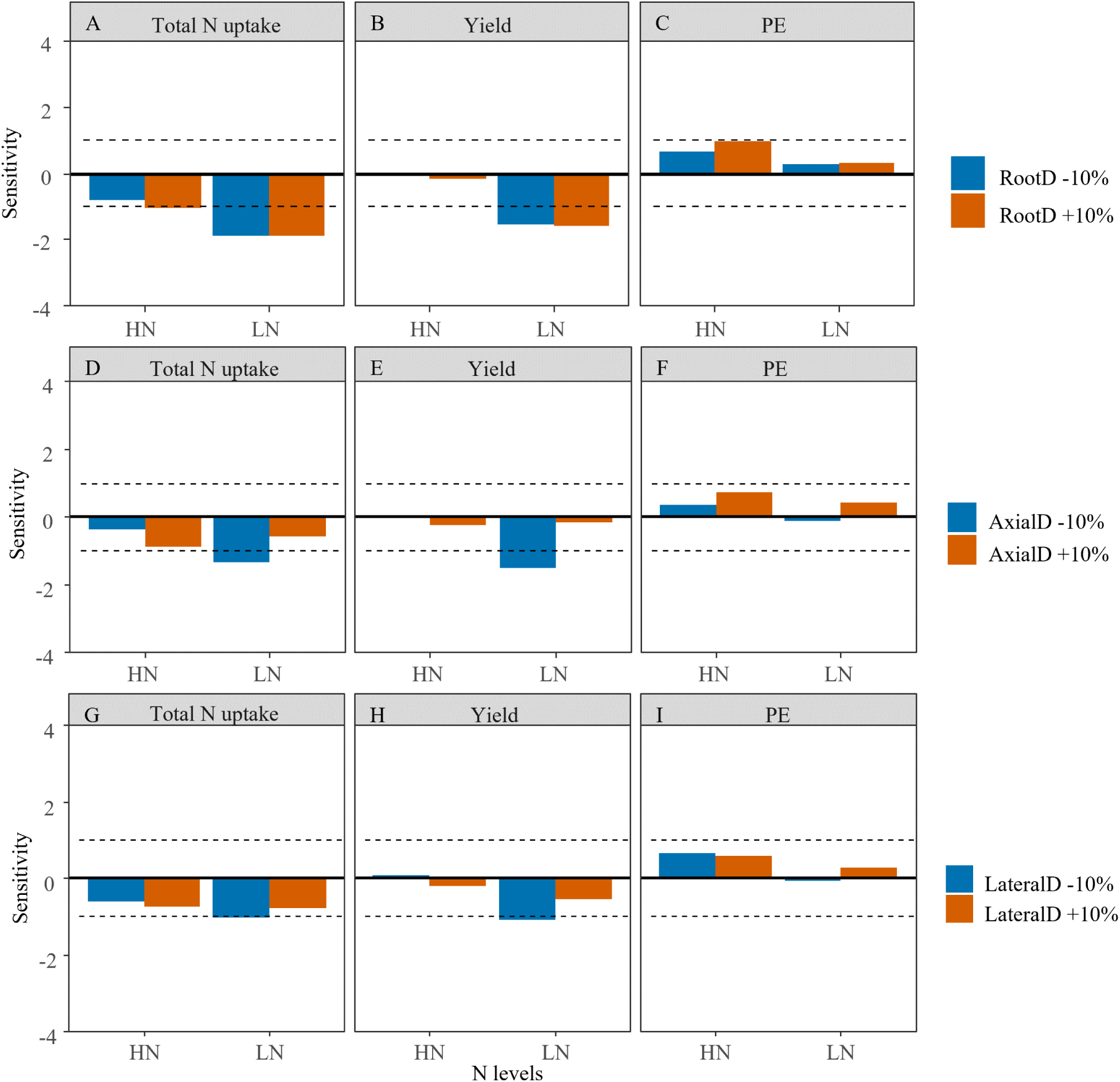
Sensitivity values of total N uptake (g/plant, A, D, G), yield (g/plant, B, E, H) and PE (g yield/g N, C, F, I) under high nitrogen condition (240kg/ha, high N) and low nitrogen condition (0kg/ha, low N) to changes in root diameter related traits: all root diameter (RootD; A, B, C), only axial root diameter (AxialD; D, E, F), or only lateral root diameter (LateralD; G, H, I). The dashed lines represent a model sensitivity of 1 and -1. Parameter and output change in the same direction give a positive sensitivity value; parameter and output change in the opposite direction give a negative sensitivity value. Blue and orange bars represent sensitivity of an output parameter to respectively a 10% decrease or increase in a trait value compared to its default.

The sensitivity of yield to the root diameter parameter values under low N was likely due to the increase in carbon costs per m root length when diameters of all root classes were increased. As these costs increased, less root length was produced leading to reduced nitrogen uptake during vegetative growth. This caused slower leaf area increase leading in turn to a lower leaf area per plant towards grain filling, which led to less carbon assimilation and allocation to the grains (Fig. 5B; S12B, D).

**Figure 5:**
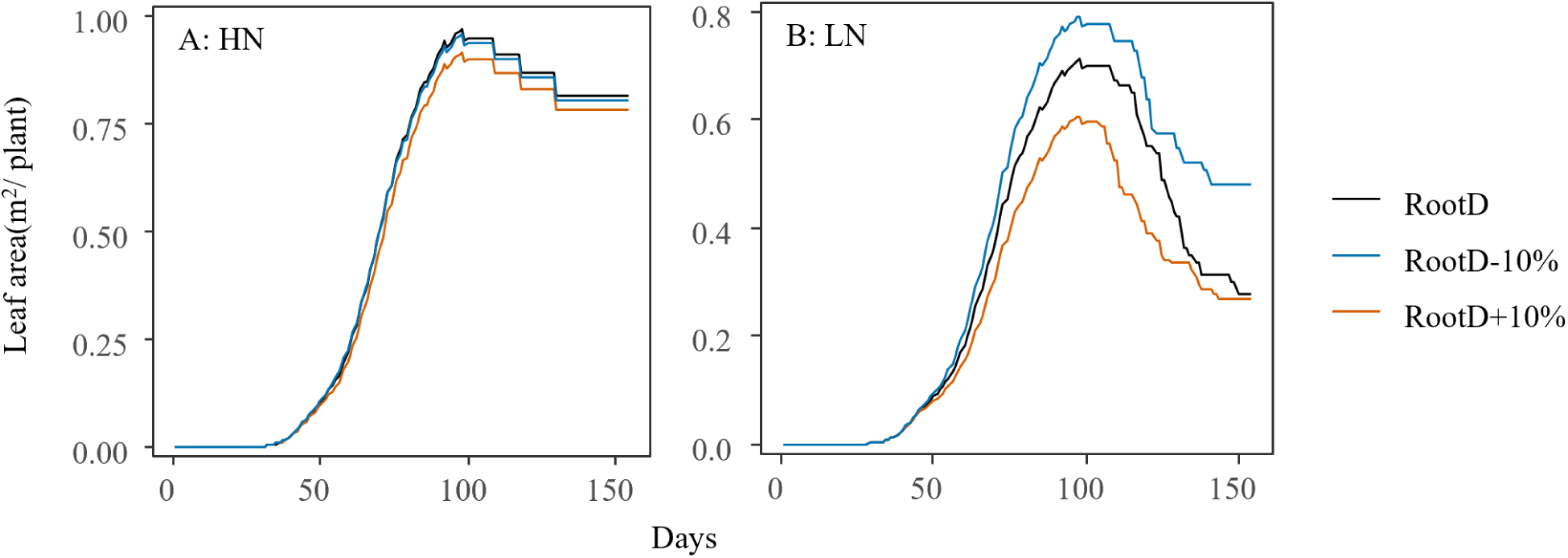
Dynamics in green leaf area per plant over the growing season when changing root diameter (RootD) under high nitrogen condition (High N, A) and low nitrogen condition (low N, B). Value ranges of y axis for high N condition is from 0 to 1 *m*^2^ and for low N condition is from 0 to 0.7 *m*^2^. Blue and orange lines represent green leaf are per plant for trait values that are respectively 10% higher of lower than their default in black lines.

Changing the maximum axial root number compared to the background genotype from 60 down to 28 had no effect on nitrogen uptake, yield and PE. However, when reducing maximum axial root number below 28, nitrogen uptake was increased especially under low N (Fig. 6A). Under low N, also yield increased with decreasing maximal axial root number, especially at numbers below 27 (Fig. 6B). Nitrogen uptake was slightly sensitive (-1.02) to the change root tissue density when increasing the value under low N (Fig. S13A). Yield and PE were not sensitive to changes in root tissue density (Fig. S13B, C).

**Figure 6:**
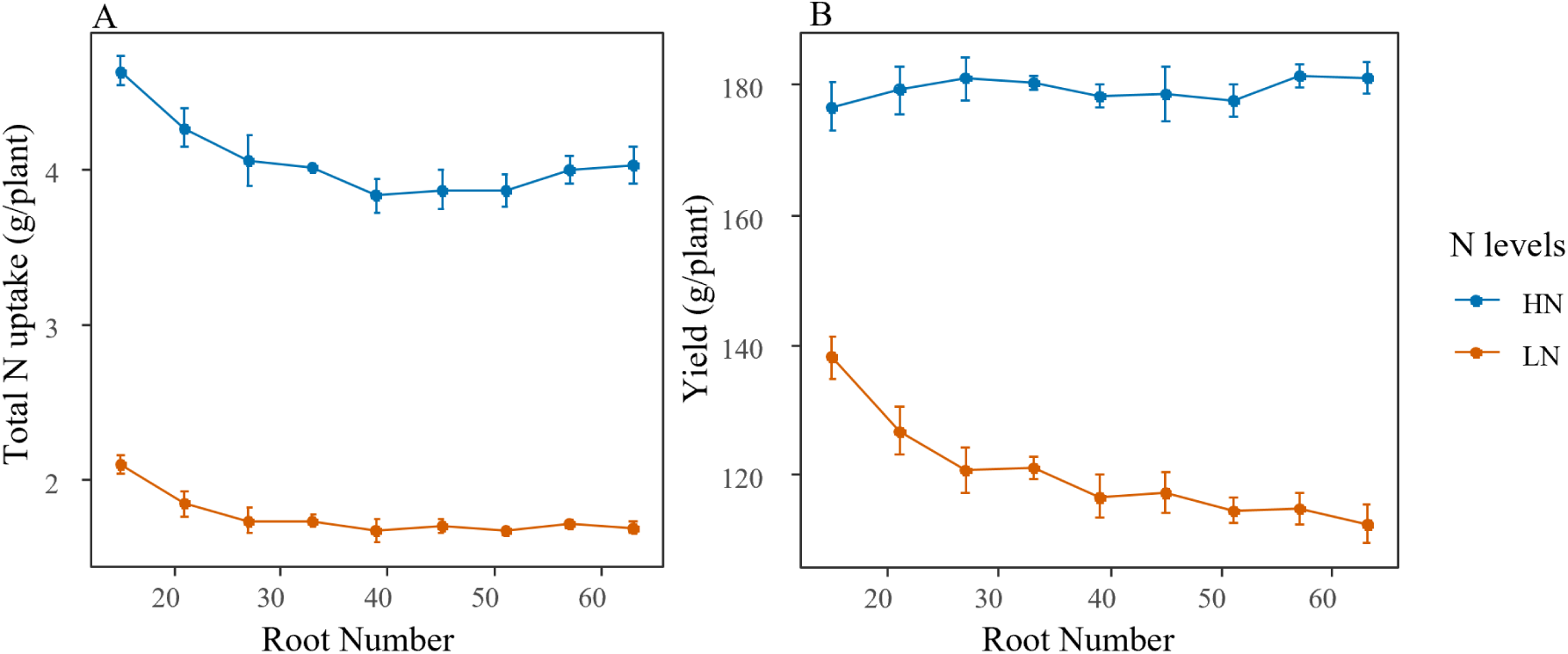
Simulated total N uptake (g/plant, A) and yield (g/plant, B) per plant across a range of maximum root numbers (*RootNum*) under both high N (blue) and low N (orange). Values represent means *±* SE (n=5).

#### Shoot traits

Nitrogen uptake, yield and PE were not sensitive to changes in leaf number with or without including an effect on the time of silking (Fig. 7A-F). Changing phyllochron changes the moment of silking without affecting leaf number; and nitrogen uptake was sensitive to changes in phyllochron under low N. This indicated that increasing phyllochron enhanced N uptake under low N. Similarly, decreasing phyllochron reduced N uptake (Fig. 7G). N uptake was sensitive to decreased phyllochron but only under high N indicating decreasing phyllochron reduced N uptake under high N (Fig. 7G). A 10% change in phyllochron, irrespective of the direction of that change, resulted in an effect on PE in the opposite direction, meaning increasing phyllochron reduced PE and decreasing phyllochron increased PE (sensitivity values ranged from -1.53 to -0.83; Fig. 7I).). Yield was not sensitive to changes in phyllochron.

**Figure 7:**
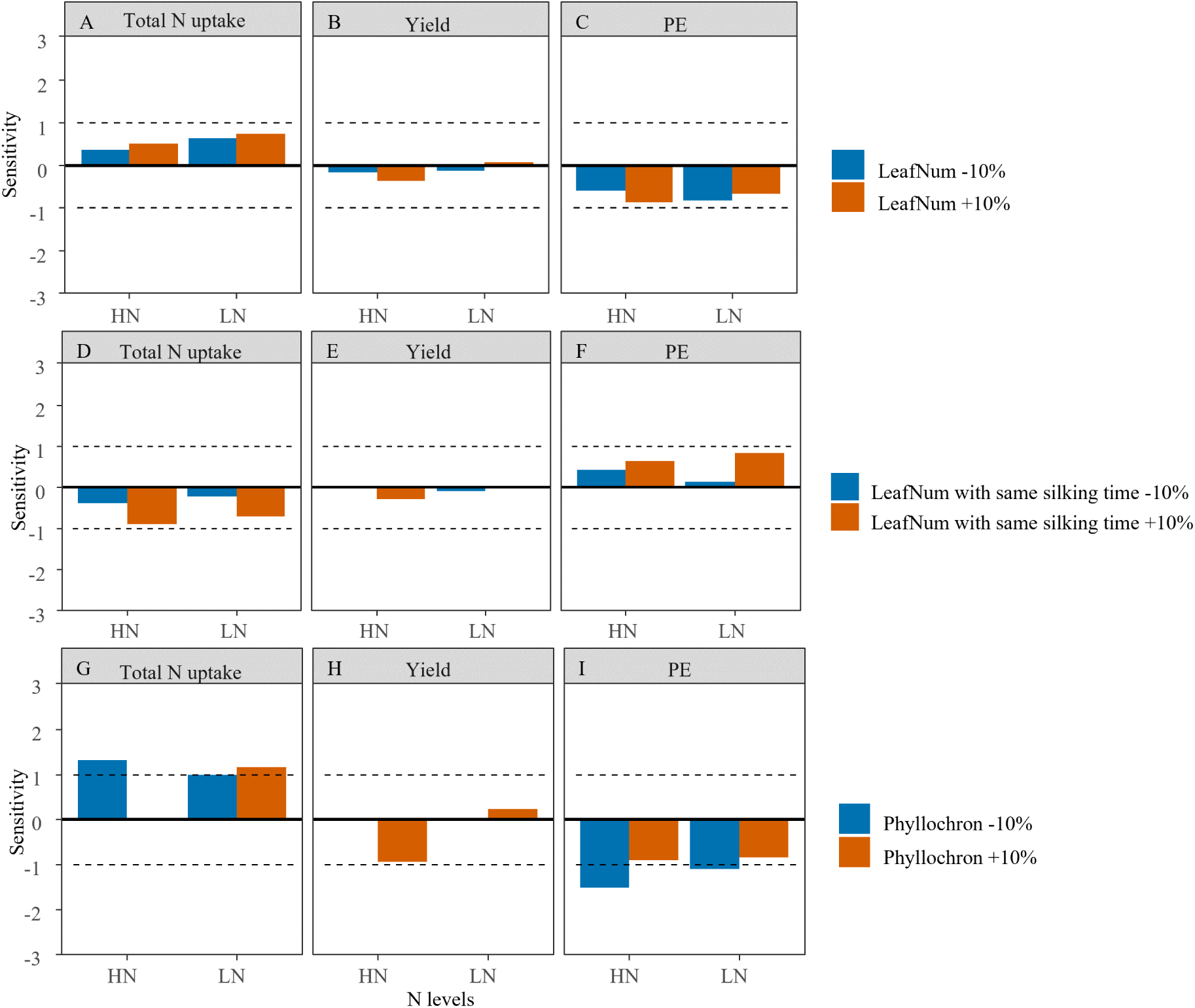
Sensitivity values of total N uptake (g/plant, A, D, G), yield (g/plant, B, E, H) and PE (g yield/g N, C, F, I) under high nitrogen application (240kg/ha, high N) and low nitrogen application (0kg/ha, low N) to changes in leaf number and time to silking related traits: leaf number (and as a consequence time of silking) (A, B, C), leaf number without affecting silking time (D, E, F), and phyllochron (and as a consequence time of silking) (G, H, I) under both high N condition and low N condition. The dashed lines represent a model sensitivity of 1 and -1. Parameter and output change in the same direction give a positive sensitivity value; parameter and output change in the opposite direction give a negative sensitivity value. Blue and orange bars represent sensitivity of the output parameter to respectively a 10% decrease or increase in a trait value compared to its default.

N uptake under high N was sensitive to decreasing the maximum photosynthetic capacity at non-limiting specific leaf nitrogen (*λ*), meaning that decreasing *λ* lowered N uptake under high N. However, under low N, nitrogen uptake, yield and PE were not sensitive (Fig. 8). Even though yield was not sensitive to *λ*, total biomass was, especially to a decrease in *λ*, and especially during early development at both N rates (Fig. 9). These positive sensitivity values indicate that increasing *λ* enhanced biomass accumulation and decreasing *λ* reduced biomass accumulation during early growth, by increasing photosynthetic capacity (*Amax*) (eq. 19). Since the effect disappeared at later growth stages, no sensitivity in yield to *λ* was observed.

**Figure 8:**
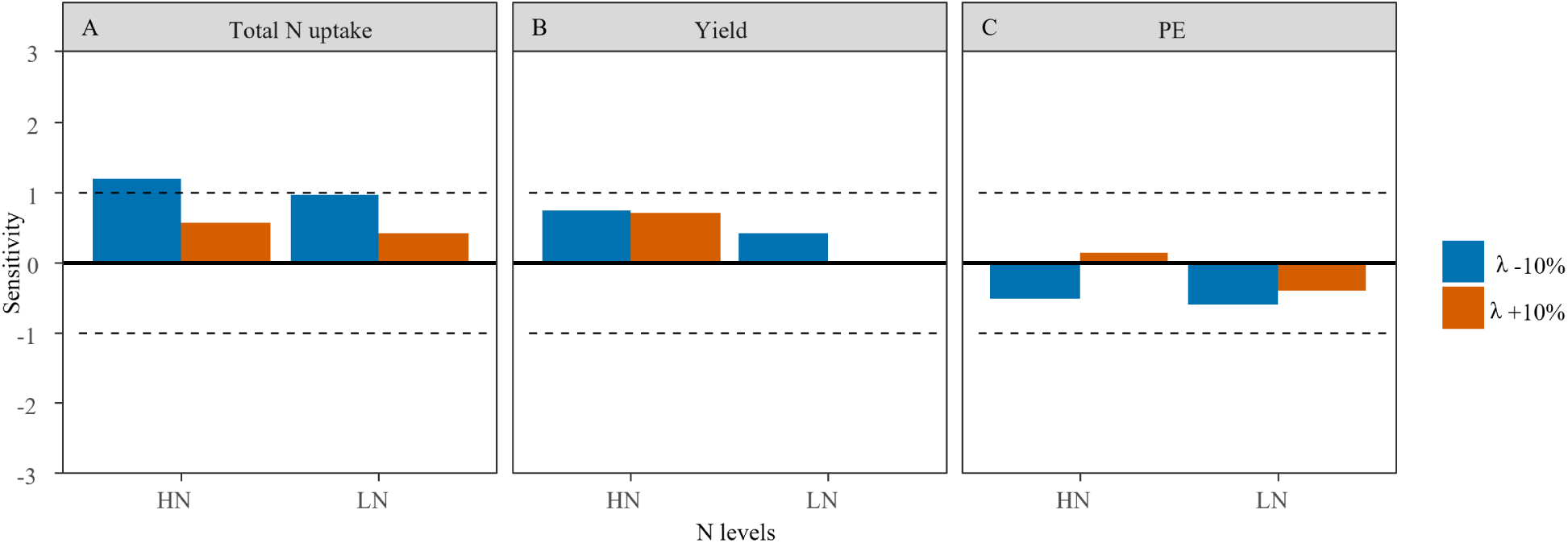
Sensitivity values of total N uptake (g/plant, A), yield (g/plant, B) and PE (g yield/g N, C) under both high nitrogen application (240kg/ha, high N) and low nitrogen application (0kg/ha, low N) to changes in the maximum photosynthesis under non-limiting leaf nitrogen *λ*. The dashed lines represent a model sensitivity of 1 and -1. Parameter and output change in the same direction give a positive sensitivity value; parameter and output change in the opposite direction give a negative sensitivity value. Blue and orange bars represent sensitivity of the output parameter to respectively a 10% decrease or increase in a traits value compared to its default .

**Figure 9:**
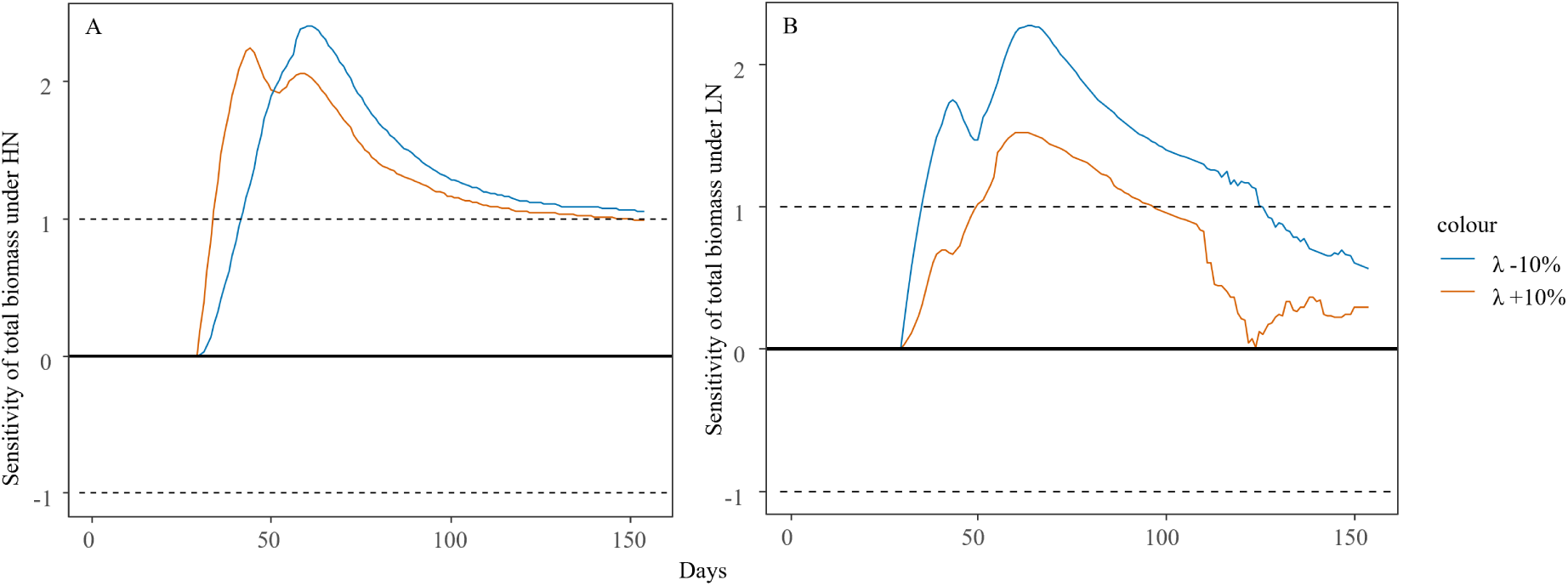
Sensitivity values of total biomass over time under both high nitrogen application (A) and low nitrogen application (B) to changes in maximum photosynthesis under non-limiting leaf nitrogen (*λ*). The dashed lines represent a model sensitivity of 1 and -1. Parameter and output change in the same direction give a positive sensitivity value; parameter and output change in the opposite direction give a negative sensitivity value. Blue and orange lines represent sensitivity of the output parameter to respectively a 10% decrease or increase in a trait value compared to its default.

#### Nitrogen reallocation

Nitrogen uptake, PE and yield were not sensitive to parameters related to grain growth rate and grain nitrogen influx rate (Fig. S14). Under both N conditions, changing potential individual grain biomass (*wmax*) by 10% resulted in sensitivities ranging from -0.346 to 0.556 for N uptake, yield and PE, while changing grain growth duration (*te*) by 10% gave sensitivities between - 0.378 to 0.209 for N uptake, yield and PE. Finally, sensitivity to changes in the N influx rate of grains (*b_g_*) ranged from -0.885 to 0.607.

## Discussion

Our model approach can capture observed differences in nitrogen uptake, yield and PE at the same magnitude for all N conditions and maize cultivars tested well enough for the purpose of testing trait effects (Fig. 2). In addition, our model reasonably well estimated the differences among cultivars even though only nine (out of more than 60) plant parameters were used to differentiate between the cultivars and this was especially true when nitrogen was not limiting. There was a significant overestimation of N uptake for cultivar ZD2 which was the oldest cultivar in the test panel, released in the 1970s. Under non-limiting nitrogen conditions, this cultivar combines a smaller root diameter with a higher specific root length compared to more recently developed cultivars such as ZD958 (Chen et al., 2014; Ning et al., 2015). The overestimation of N uptake for ZD2 indicates that additional processes may have to be included in the current model to improve the representation of old cultivars. A possible difference between old and more recent cultivars is in their root anatomical characteristics. These may interact with soil chemical and physical characteristics through, for instance, penetration strength of roots, which in turn impacts N uptake. The differences between observed and simulated N uptake may thus related to the trade-off between penetration strength and soil N forage of thin roots. Soil texture and root penetration strength were not part of this study. To capture plant architectural, developmental and physiological genotype-specific traits related to PE, our model is a suitable tool.

### Maize traits related to N uptake and PE

#### The role of root architectural and physiological traits in N uptake and PE

Our simulations suggested that root traits such as root diameter are more relevant to N uptake and PE than physiological traits related to root N uptake mechanisms, especially under low N conditions. Differences in *I_max_* have been reported between different regions along roots from highest at root tips to lowest at the root base, as well as between axial or lateral roots (Lazof et al., 1992; Sorgona et al., 2011). However, such observations were typically made in experiments at the scale of individual root segments, and usually over a short time span. Differences in *I_max_* were also identified among maize genotypes at the root system scale, but these were found only weakly correlated to nitrogen uptake rate at plant level (Pace and McClure, 1986). This is also the conclusion from the current modelling exercise.

In an earlier modelling study (Dunbabin et al., 2004) plasticity in ion-uptake mechanisms was more relevant for N uptake in herringbone type root systems typical for dicots, than in dichotomous type root systems such as in maize. Similarly, within the range of measured *I_max_* for different maize root classes, shoot biomass was not sensitive to *I_max_* over soil nitrate ranging from 20kg/ha to 200kg/ha (York et al., 2016). In general, the capacity for nitrogen uptake plays a more important role when competition or compensation for N between roots of a same plant is low, due to low root density. In addition, a lower root length density in the top soil layer (0-20 cm) was found with higher root system efficiency of N for modern maize cultivars (released after 1990s) compared with old maize cultivars (Chen et al., 2014). Combining with our simulation results (Fig. 3), all together indicated the competition for soil N among root segments with a plant plays a more important role than capacity of N uptake for individual root segment. Since maize plants usually have dense root systems, it would be less important to consider capacity for nitrogen uptake such as *I_max_* as a primary target trait in a maize breeding program aimed at high nitrogen use maize genotypes. In contrast, breeding for traits allowing better soil exploration will be very relevant.

Root diameter inheritance in our model is based on the principles of the ArchiSimple model (Pagès et al., 2014). Therefore, decreasing root diameter caused carbon investment to go into additional root length, extending the root system into previously unexplored soil allowing additional N uptake. This reduced competition among root segments for limited soil nitrogen within each soil cell, through a reduction of root surface area per cell. In *in vivo* studies, longer root hairs can increase nitrogen uptake under low N which also means roots with small diameter can increase their N depletion zone (Saengwilai et al., 2021). In line with our findings, a smaller axial root diameter was found to provide an adaptation to low N conditions (Gao et al., 2015). In addition, quantitative trait loci for smaller root diameter were associated with higher N uptake and finally higher yield (Coque et al., 2008). However, since primary root diameter can be correlated with anatomic traits which affect penetration strength, N use efficient genotypes generally have larger root diameters than N use inefficient genotypes under low nitrogen in field conditions (Yang et al., 2019) which is contradicting our results. In addition, the diameter of basal roots were positively correlated with root length under field conditions indicating the importance of penetration strength (Wu et al., 2016). Both of the examples hint at a trade-off between root expansion and root penetration strength not included in our current model. Further work on the role of root diameter on N use should therefore consider competition for resources between plants in relation to soil compactness and related requirements for penetration strength of roots. Alternatively breeding towards plants with a lower primary root number (Fig. 6) and lower root tissue density (*RTD*) could lead to reduced carbon costs at equal penetration strength allowing for root system expansion.

#### Trade-off between biomass accumulation and yield

Phyllochron determines the rate of creation of new sinks for carbon and nitrogen. Together with final leaf number, phyllochron determines the duration of vegetative growth, which controls the carbon sink:source ratio during plant growth. Increasing phyllochron led to individual leaf area of lower leaves becoming larger, since fewer sinks competed for resources in the early growth stage. In line with our findings, a positive correlation was found between phyllochron and leaf size in rice (Rebolledo et al., 2012; Miyoshi et al., 2004). Based on our model assumption that root sink strength is a fraction of leaf sink strength, increasing both traits improved root growth. However, both phyllochron and final leaf number hardly affected simulated yield. Increasing phyllochron can increase leaf area during vegetative growth, which meant a larger assimilate requirement to develop vegetative structures. Also, the final grain growth is controlled by both grain sink strength and available assimilates. Therefore, an increase in phyllochron ended up with no significant increase in yield. Similarly, increasing leaf number also hardly affected yield since yield formation did not only depend on source assimilation, but also on grain sink strength.

Yield and PE were not sensitive to leaf photosynthetic capacity related to leaf N while total biomass was sensitive only at early growth stages (Fig. 8B, C, Fig. 9). The combination of a reduction in leaf N sink strength with developmental stage with changes of biomass accumulation at early growth stages, may have led to limited changes in photosynthetic capacity. Sink-related traits have been reported to play an important role in limiting both plant growth and grain development in annual crops (Borrás et al., 2004; Serrago et al., 2013; Long et al., 2006). However, we observed that simply increasing grain sink strength, by increasing potential grain growth rate (*wmax* and *te*), also hardly affected final yield. Thus, sink and source rather co-determine final yield, and only accounting for the function of just the sink for or the source of grain carbon hardly improves yield.

#### Breeding for an increase in PE

PE (kg grain/ kg N in the plant) is a complex trait that is affected by the nutrients present in the plant, it distribution among organs, photosynthetic traits and sink potentials. Component traits like photosynthesis related traits can have higher heritability than compound traits such as yield, can have stronger genetic correlation and are easier to measure across a wide range of genotypes during early plant growth (Gallais and Coque, 2005). Therefore, a FSP model like ours is a valuable tool to dissect PE into plant architectural and functional traits like photosynthetic capacity, and to identify which of these could be future targets in breeding programs.

In order to breed for a more nutrient use efficient maize, root morphological traits such as root diameter are potentially interesting breeding targets. The root system architecture is directly associated with biomass accumulation and grain yield for maize (Hammer et al., 2009). Genetic variation in root diameter has been reported (Yang et al., 2019; Coque et al., 2008) giving scope for breeding. However, there are still gaps in our understanding of trade-offs as thinner roots have the potential to explore a larger soil volume by increasing root length while it may equally reduce its penetration strength into more compacted soil. Phyllochron and the maximum photosynthesis under non-limiting leaf nitrogen (*λ*, eq. 23) also appeared to be interesting plant traits. Even if phyllochron seems to be of minor importance for final yield, this trait determines vegetative biomass accumulation during early growth and thus N uptake of the plant. Therefore, breeding for higher values of this traits can potentially improve nitrogen uptake of plants since larger leaf area positively correlates with root biomass especially under N-limiting condition. Genetic variation exists for the phyllochron, and this is not always associated with final leaf number (Lacube et al., 2020; Padilla and Otegui, 2005). Therefore, it is possible to breed for this traits independent from leaf number. Similar to phyllochron, *λ* can be an interesting trait to breed for to increase N uptake of maize, through enhancing biomass accumulation. Genes identified for photosynthetic capacity in maize directly resulted in changes in biomass accumulation and N uptake (Li et al., 2020; Wang and Li, 2019). Our model indicates plant traits that seem important based on our understanding of whole plant growth and development, may help to target breeding for nitrogen use efficiency. However, combining results from our model and previous studies, we would argue that simply improving a limited number of plant traits cannot result in a significant increase in grain yield. For such an increase likely several plant traits will simultaneously have to be changed. For instance, yield was not found sensitive to either grain sink strength or *λ*.

### Potential next steps in modelling

Plastic responses of plant traits to environmental conditions play an important role in root growth and development and thus root system architecture (Schneider, 2022). Our current approach only included a limited number of generic plastic responses to nitrogen like making the allocation of produced assimilates to roots or shoot dependent on leaf N concentration and making the N uptake rate (both the low and high affinity transport) dependent on plant N concentration. However, a number of potentially also relevant plastic reactions in response to plant and soil nitrogen queues have not been included in our approach, due to a lack of suitable empirical data to account for genotype-specific difference in plasticity. To prioritize plasticity by its importance on plant performance for further study, results of sensitivity analysis can be used. For instance, based on results of the sensitivity analysis (Fig. 4, 6), changes in root diameter and root number were significant in plant N uptake and yield, especially when N application was reduced. Differences in these traits between various N conditions within a same genotype have also been observed through experiments (Saengwilai et al., 2014a; Schneider et al., 2021). Both *in silico* and *in vivo* results together indicates the importance of plasticity in these two traits on plant N use and is worth further study. Therefore, for future studies with FSP models, plastic responses such as in root diameter can be considered for integration to identify and quantify effects of variation available among genotypes. After implementing the relevant plant mechanisms, conditions like daily temperature, radiation and soil N dynamics can then be considered as input to further predict plant performance more precisely. If simulations would show potential for improved nitrogen uptake or internal use efficiency these genotype-specific parameters could then be considered relevant target traits for breeding towards more efficient N usage maize across environments.

## Acknowledgement

The authors are grateful to Xiaochao Chen, for sharing original field datasets. The authors declare no conflicts of interest.

## Author Contribution

JL, JBE, TJS and LY designed the research; JL, JBE and TJS developed the model and analysed data; JL wrote the paper. JBE, TJS, LY and GM revised the manuscript and contributed to the finalization of the manuscript.

