## Supplementary Figures and Tables for "Identifying and quantifying the contribution of maize plant traits to nitrogen uptake and use through plant modelling"

### Supplementary materials

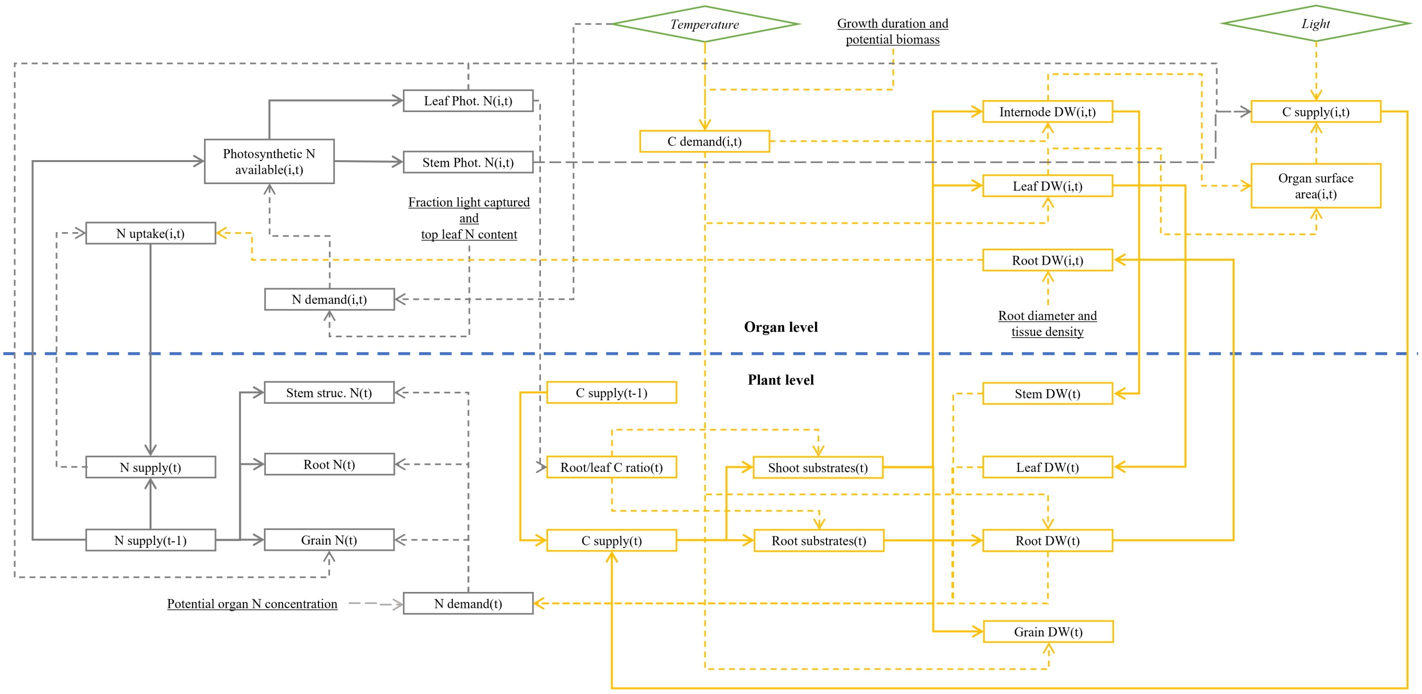


Supplementary Figure 1: Flowchart of the nitrogen (N, grey) and carbon (C, yellow) flows and related processes in the model. Solid arrow lines indicate matter flows determining N and C accumulation at each simulation step (per day). Dotted arrow lines represent information flows. The rectangles represent state variables. The input parameters are underlined. The diamonds represent environmental inputs. The top half refers to organ level flows and states represented by individual leaf, internode and root segments. The bottom half refers to plant level flows and states calculated from the addition of individual leaves, internodes or root segments into whole plant organ types (leaves, stem or root). In addition, grain is treated as separate organ type (individual grains are not simulated so not represented).

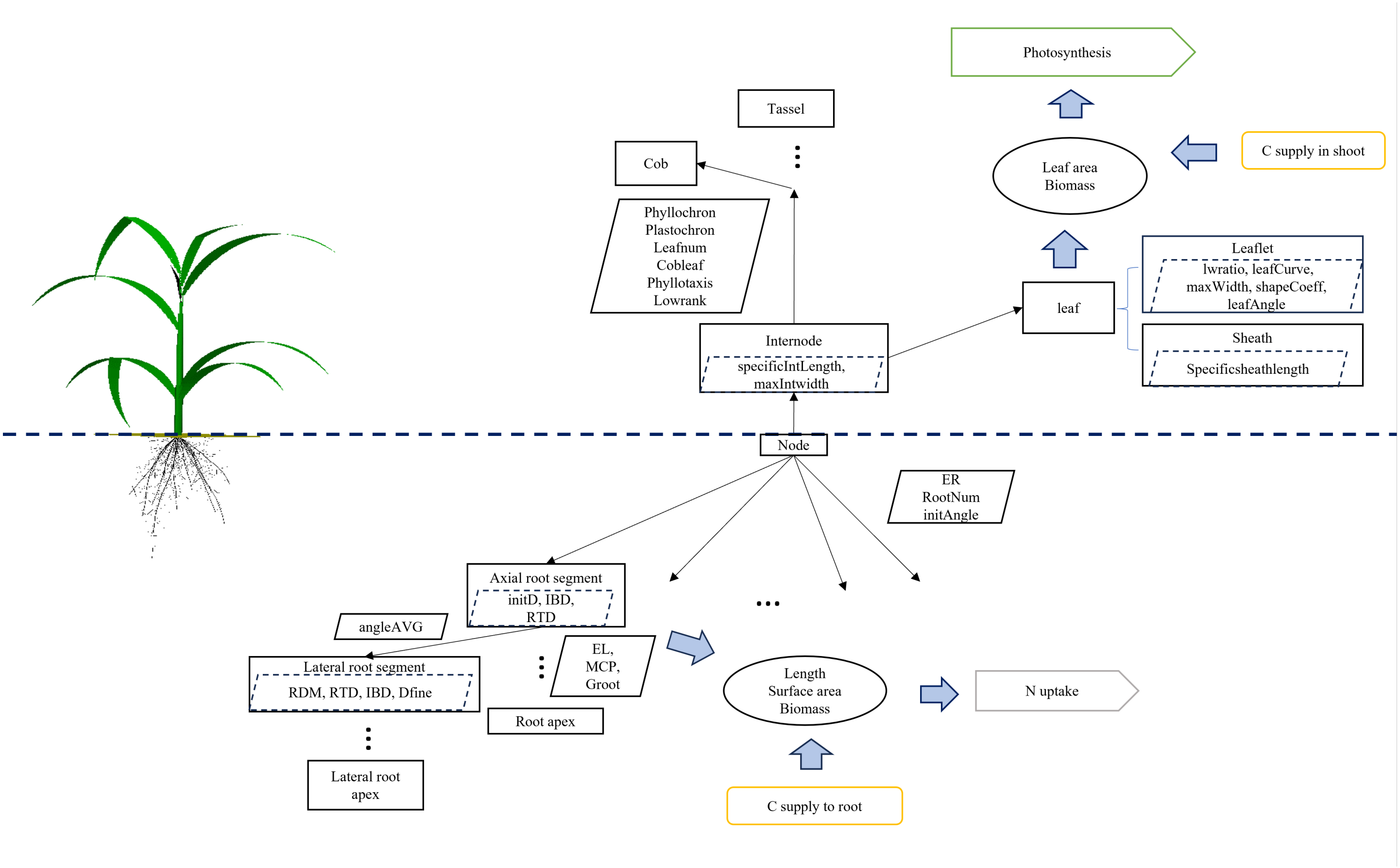


Supplementary Figure 2: Illustration of geometry for the maize FSP model (left) and how the model defines this (right). Rectangles represents plant organs with visualized geometric shapes. Parallelograms represent the geometric parameters defining the potential organ shapes. Parallelograms with solid borders represent global parameters affecting the plant architecture while the parallelograms with broken borders represent parameters directly influencing specific organ shapes. The rectangles in yellow with smoothed edges represent available carbon to roots and shoot. For more detailed information on the carbon and nitrogen flows see Fig. S1. The ovals represent the major outputs determined by both available carbon and organ shapes and these were used to calculate photosynthesis and N uptake.

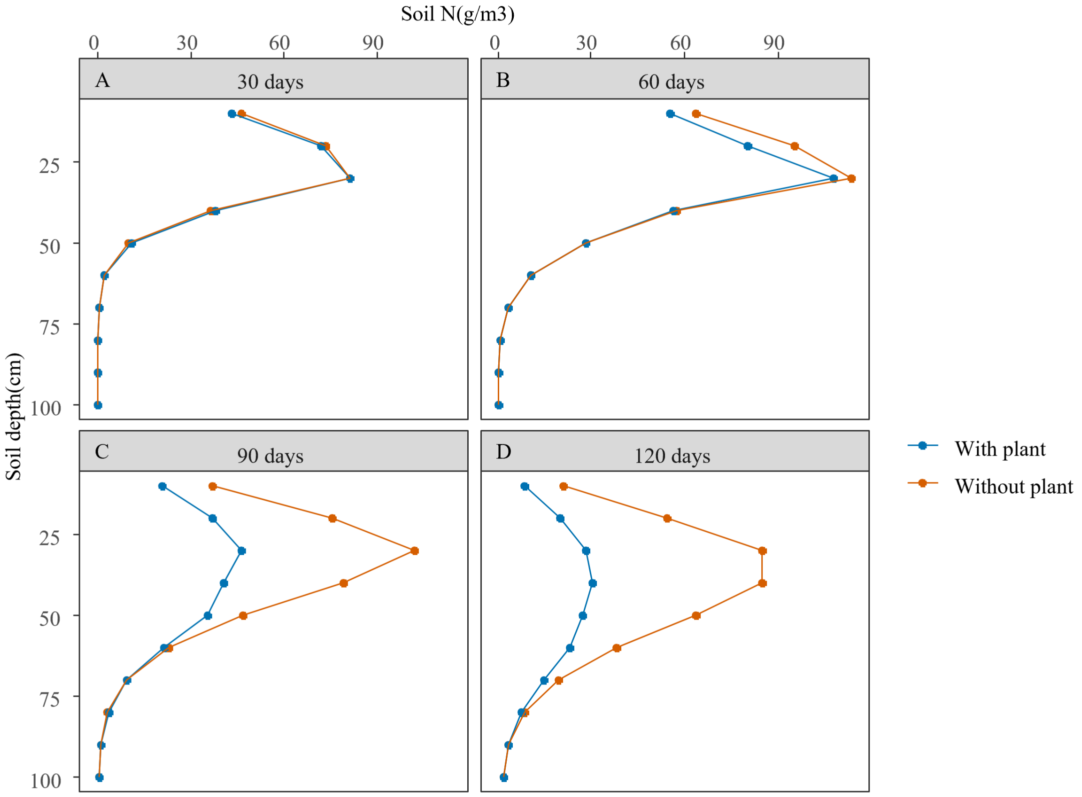


Supplementary Figure 3: Simulated soil N distribution during plant development under high N condition (5.714 $\mu$mol/m$^3$). Blue color represents soil N distribution in soil with a growing plant. Orange color represents soil N distribution in soil without a plant. Different panels represents different days after application of N fertilizer.

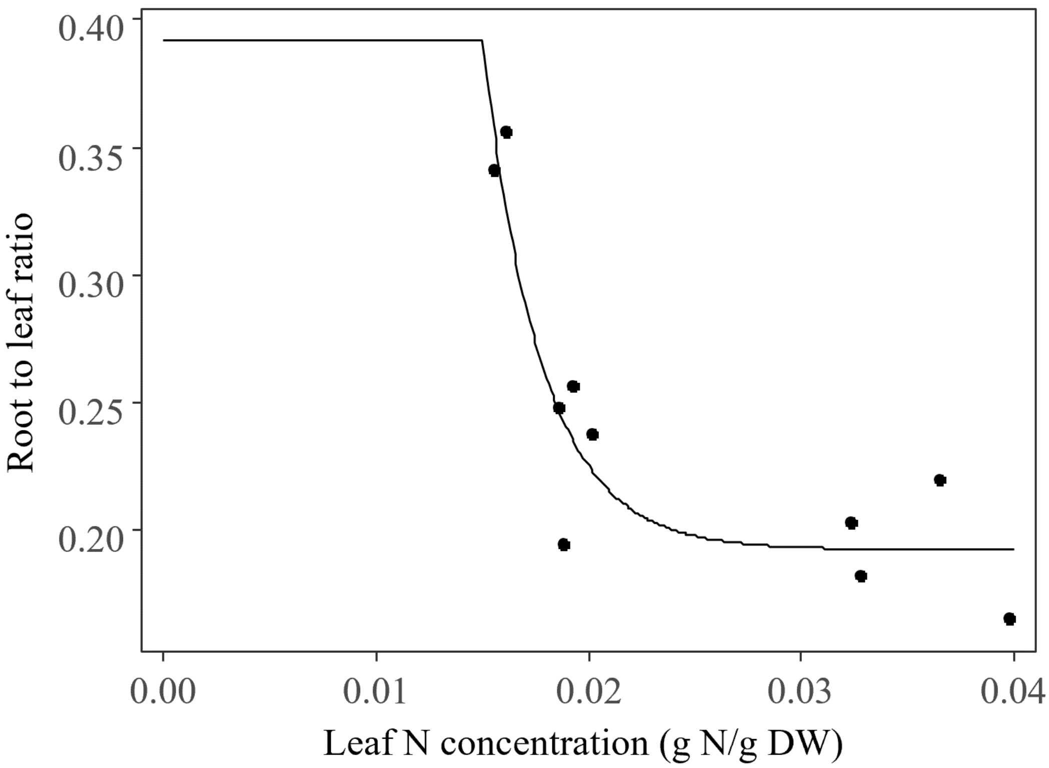


Supplementary Figure 4: Root to leaf ratio in response to leaf N concentration. The dots represents experiment data for cultivars ZD958 and XY335 and the solid line represents the fitted relationship implemented into the model.

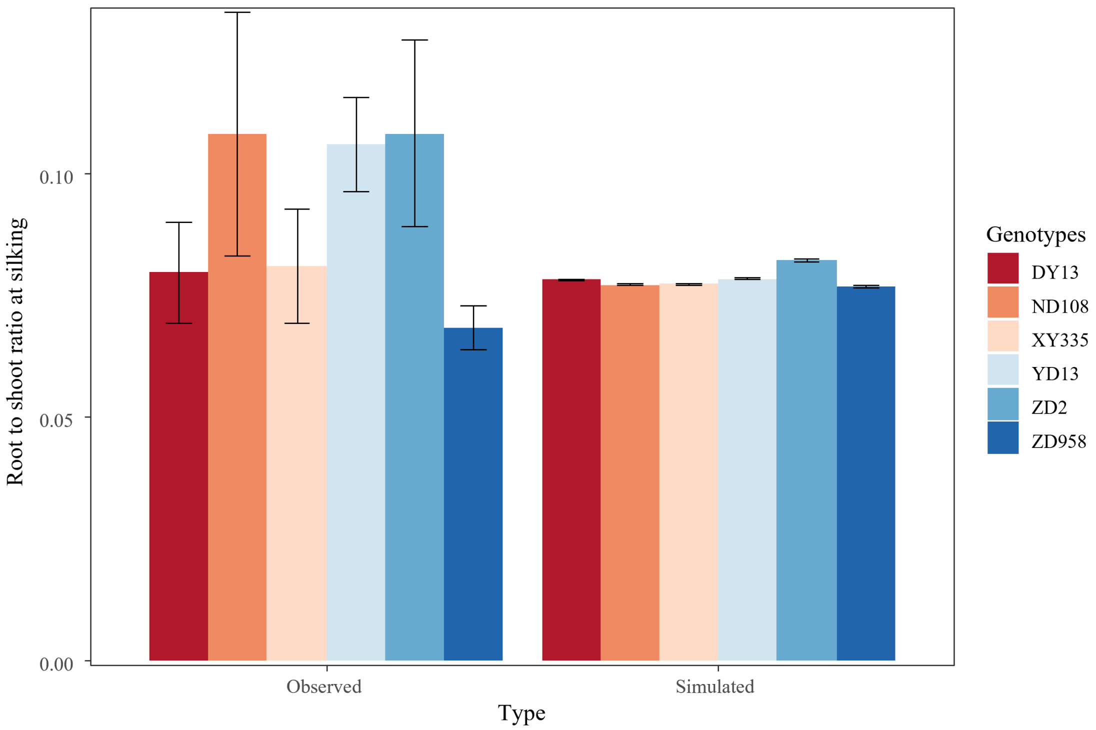


Supplementary Figure 5: The evaluation of root to shoot ratio. The observed root to shoot ratios of each maize genotype were derived from the field experiment under high N condition (240 $kg/ha$ and 5.714 $\mu$mol/m$^3$) in 2010 from Chen et al. (2013). The simulated root to shoot ratios were the output based on 2010 light and temperature data under high N condition. The Different colors represented different genotypes. The values are means $\pm$ SE ($n_{obs}$=4 and $n_{sim}$ = 5. )

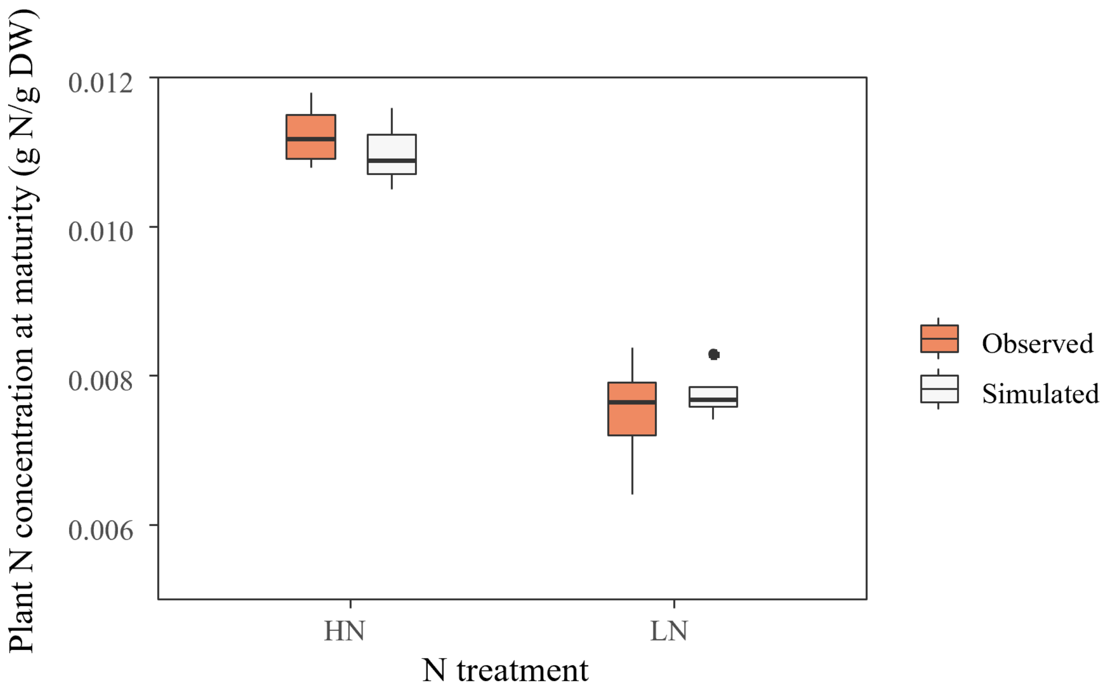


Supplementary Figure 6: Boxplot of plant tissue N concentration at physiological maturity. The default cultivar used here was ZD958 in order to illustrate plant N status under limiting and non-limiting N conditions.The orange boxes represented the plant N concentration of ZD958 in 2010 from Chen et al. (2013). The white boxes represented simulated plant N concentration of ZD958 based on the 2010 light and temperature data. HN represents high N level (5.714 $\mu$mol/m$^3$) and LN represents low N level (0 $\mu$mol/m$^3$). n for observed data is 4 and for simulation data is 5.

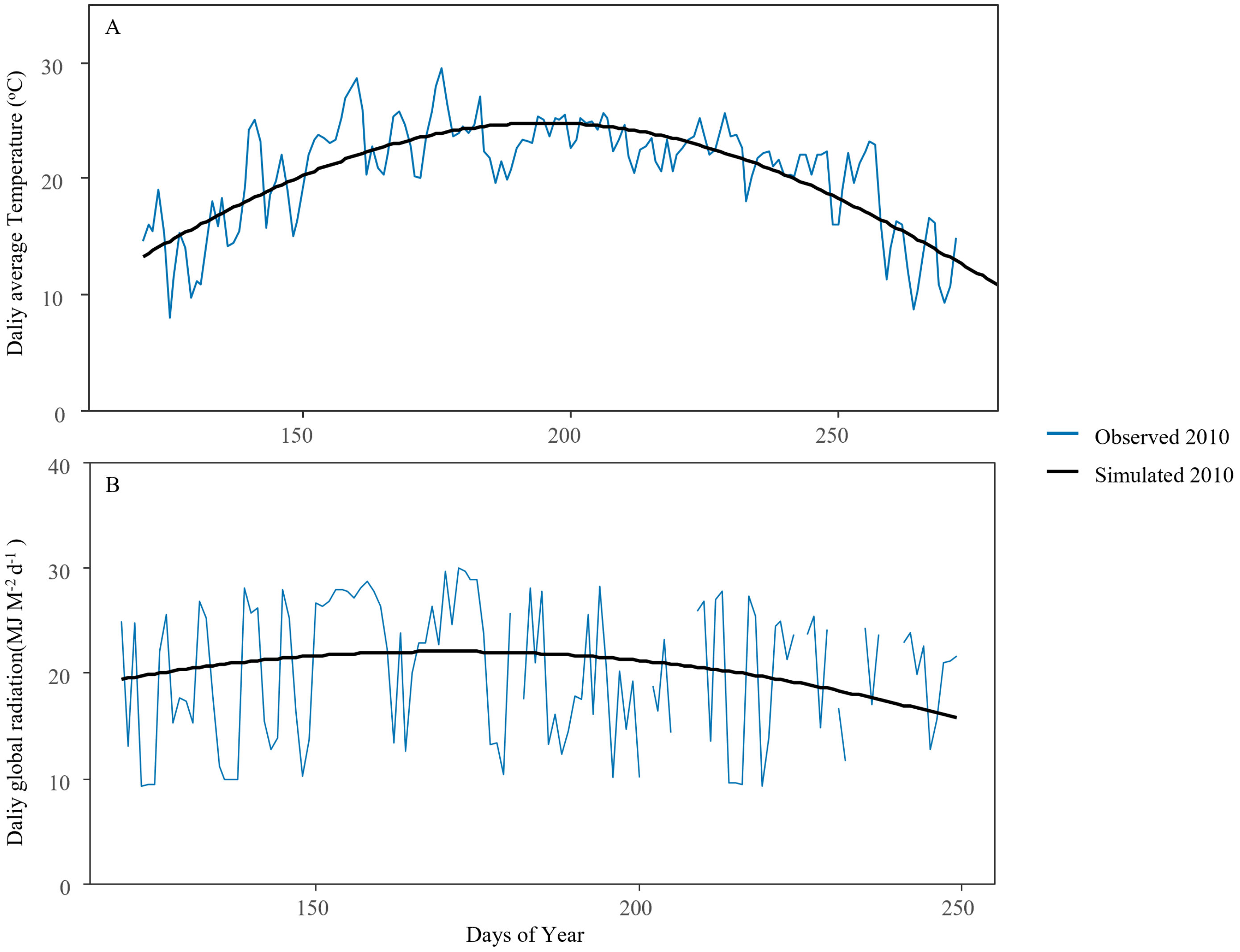


Supplementary Figure 7: Daily average temperature (A) and daily global radiation (B) for the maize growing season in Lishu, Jilin in 2010 (Chen et al., 2013). Blue lines link actual measured data for 2010, where breaks represent days with missing values. Black lines represent the smoothed data for 2010.

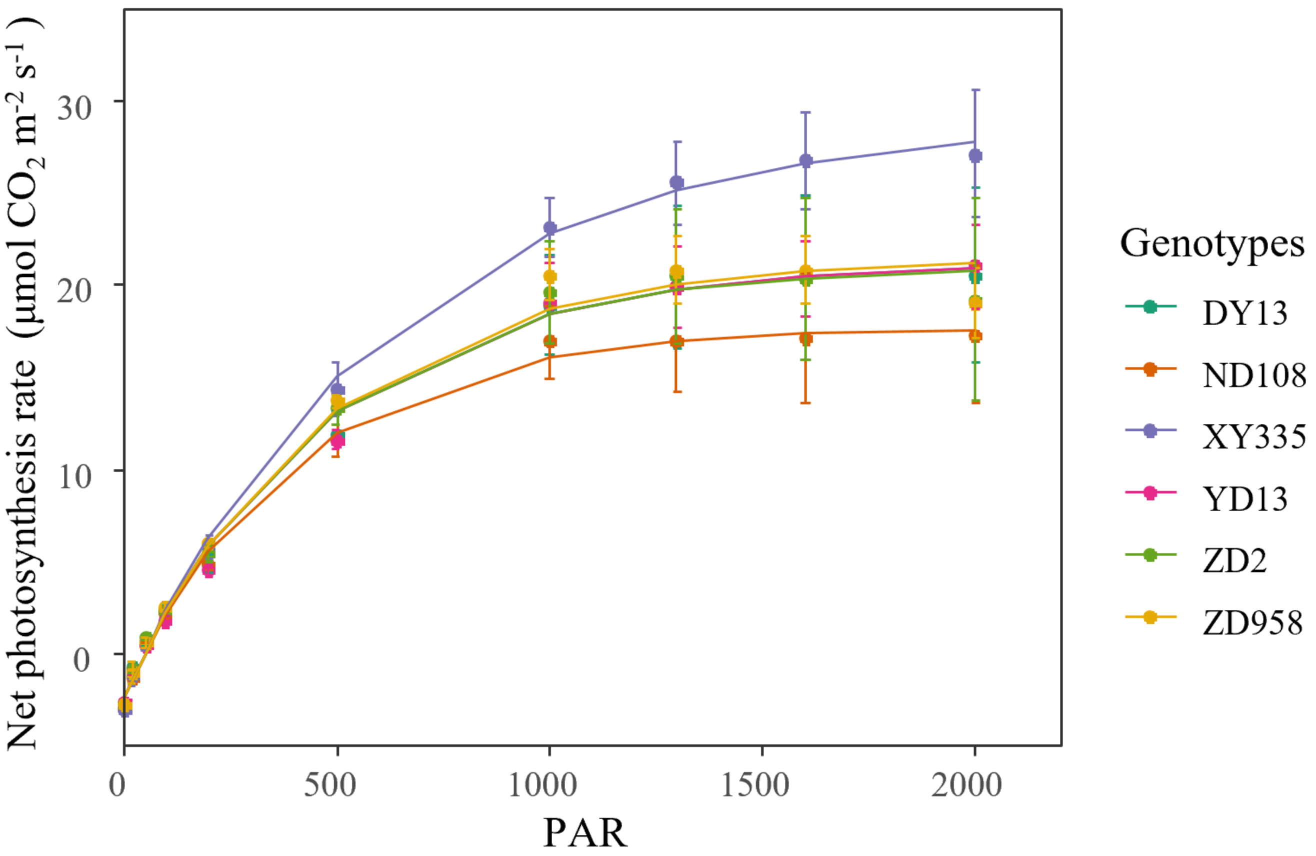


Supplementary Figure 8: Leaf photosynthesis measurements from plants grown in the field under high nitrogen condition during the 2011 growing season for six maize cultivars (points) and fitted light response curves for non-limiting leaf nitrogen (eq. 23), where the maximum net photosynthesis rate $\lambda$ was considered a cultivar specific parameter. Observed values represent means $\pm$ SE (n=4), based on the data from Chen et al. (2014).

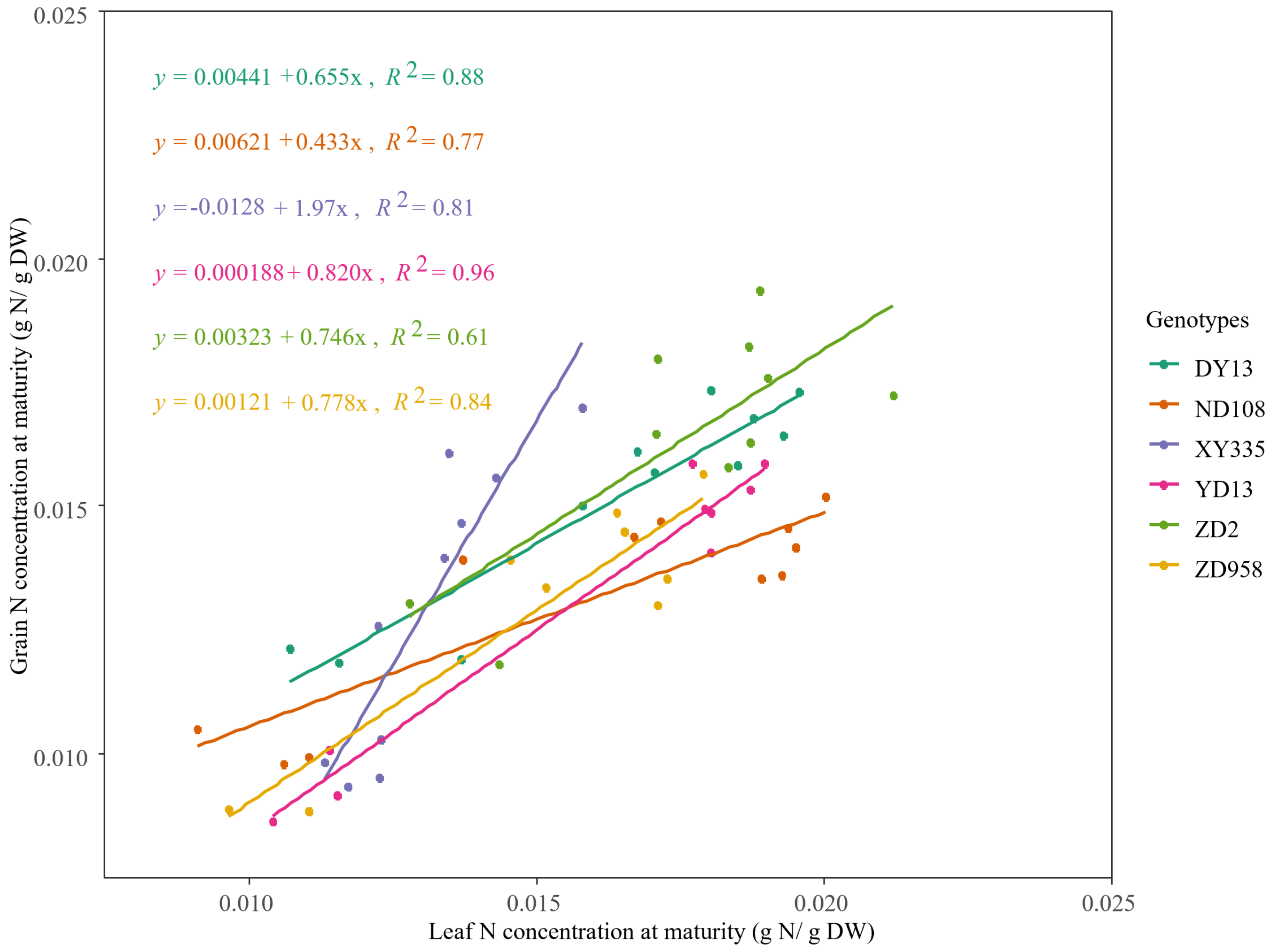


Supplementary Figure 9: Relations between grain N concentration and leaf N concentration as fitted on observed data of six maize cultivars collected at physiological maturity during the 2011 growing season (original data from Chen et al. (2014).

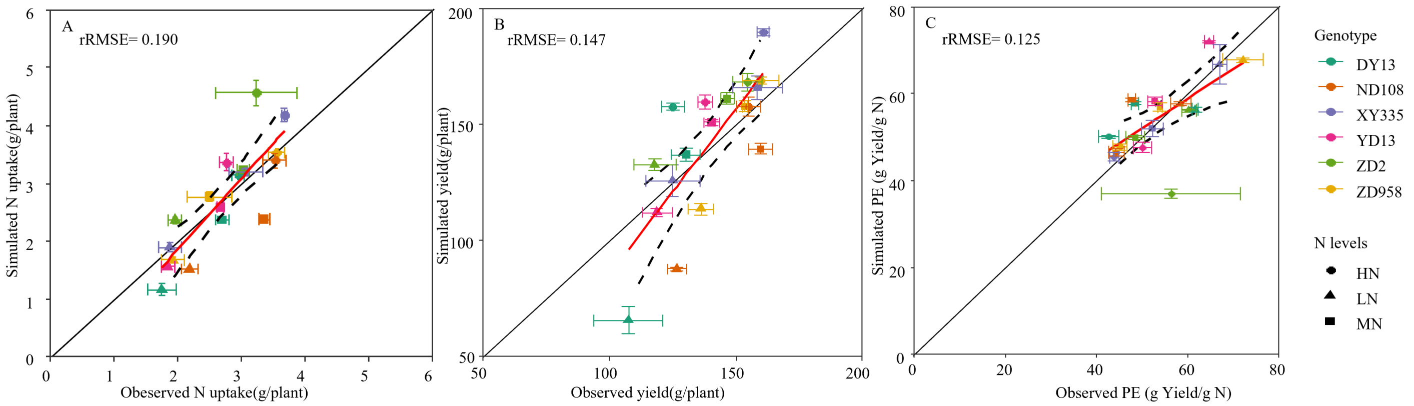


Supplementary Figure 10: Scatter graphs of simulated versus observed N uptake (A), yield (B) and PE (C) for six maize cultivars grown at high (240kg N/ha, HN), medium (120kg N/ha, MN) and low (0kg N/ha, LN) N fertilizer application rates. Observed daily average temperature and radiation were used in the simulations. Observed data are from experiments in 2010 reported in Chen et al. (2014), where points and error bars represent means ± SE (n=4). The solid lines represent the fitted line between observed and simulated data while the 95% confidence intervals of the fitted lines is represented by the area between the two broken curves.

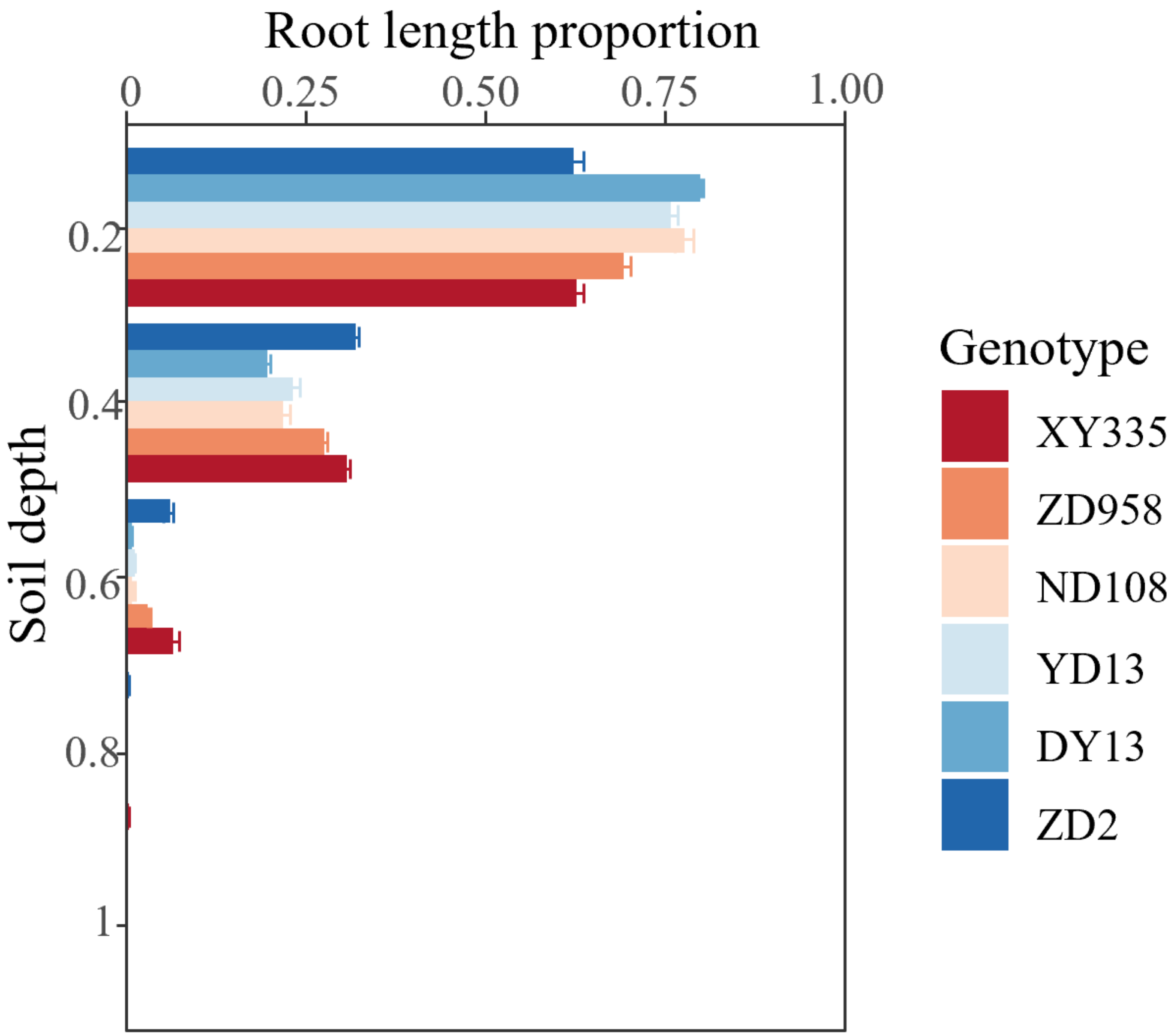


Supplementary Figure 11: Simulated root length proportion for the six maize genotypes. The simulations were run using observed weather condition in 2010 at high N (240 kg/ha) fertilizer application rates. The bars and error bars represent means ± SE (n=5).

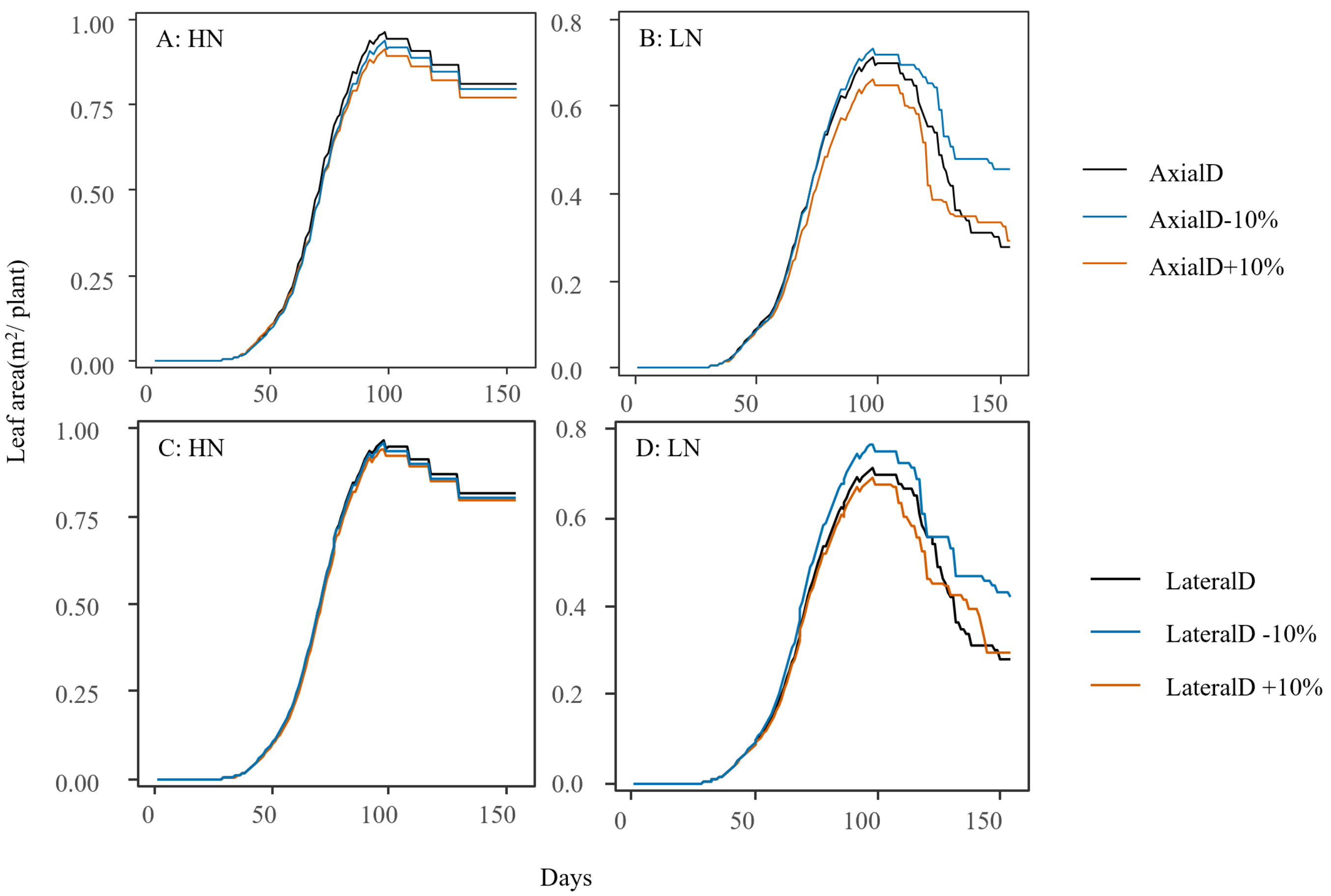


Supplementary Figure 12: Dynamics in green leaf area per plant over the growing season when changing root diameter related traits: only axial root diameter (AxialD, A, B), or only lateral root diameter (LateralD, C, D) under high nitrogen condition (High N, A, C) and low nitrogen condition (low N, B, D). Value ranges of y axis for high N condition is from 0 to 1 $m^2$ and for low N condition is from 0 to 0.7 $m^2$. Blue and orange lines represent green leaf are per plant for trait values that are respectively 10% higher of lower than their default in black lines.

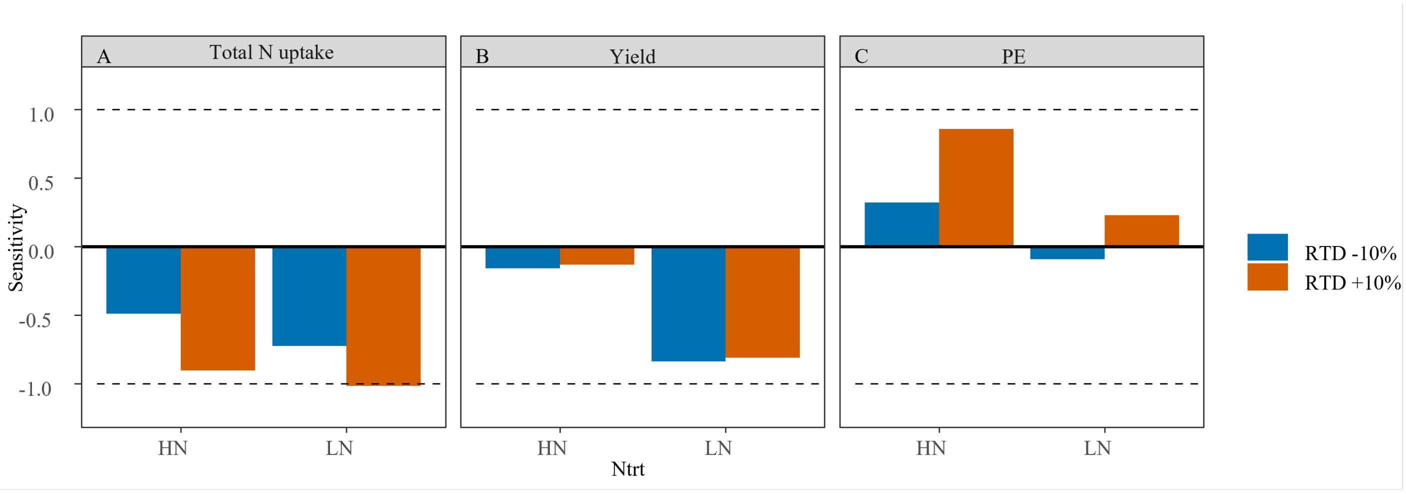


Supplementary Figure 13: Sensitivity values of total N uptake (g/plant, A), yield (g/plant, B) and PE (g yield/g N, C) under high nitrogen condition (240kg/ha, high N) and low nitrogen condition (0kg/ha, low N) to changes in root tissue density. The dashed lines represent a model sensitivity of 1 and -1. Parameter and output change in the same direction give a positive sensitivity value; parameter and output change in the opposite direction give a negative sensitivity value. Blue and orange bars represent sensitivity of an output parameter to respectively a 10% decrease or increase in a trait value compared to its default.

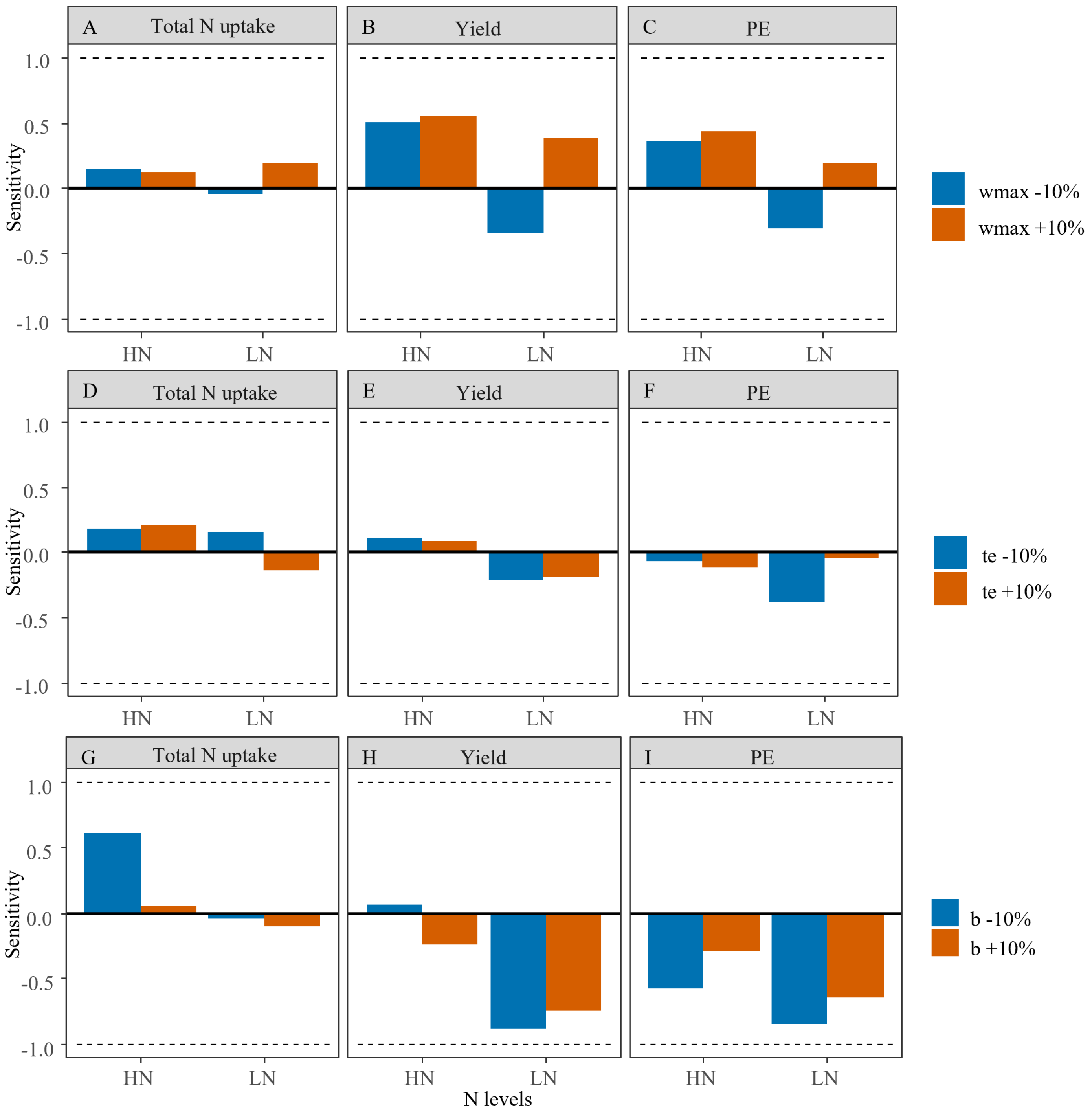


Supplementary Figure 14: Sensitivity values of total N uptake (g/plant, A, D, G), yield (g/plant, B, E, H) and PE (g yield/g N, C, F, I) under high N condition (240kg/ha, high N) and low N condition (0kg/ha, low N) for nitrogen remobilization related traits: potential grain dry weight (wmax, A, B, C), grain growth duration (te, D, E, F), and grain N influx rate ($b_g$, G, H, I). The dashed lines represent a model sensitivity of 1 and -1. Parameter and output change in the same direction give a positive sensitivity value; parameter and output change in the opposite direction give a negative sensitivity value. Blue and orange bars represent sensitivity of the output parameter to respectively a 10% decrease or increase in a trait value compared to its default.

Supplementary Table 1: List of management and environment parameter values used for model validation (2010). The parameter values were derived or fitted from experimental and local weather station data.

| Parameter | Description | 2010 |
|---|---|---|
| startingdate | Sowing date | 129 |
| delay | Time lag between sowing and emergence | 17 |
| DNapp2 | Second N application at V8 | 60 |
| harvest date | Growing days after plant emergence | 138 |
| rowDistance | Distance between two rows (cm) | 60 |
| plantDistance | Distance between two plants (cm) | 30 |
| a | Year average temperature | 9.121 |
| b | Seasonal variation in daily average temperature | 15.72 |
| c | Day of the year when temperature is the yearly average temperature | 104 |
| Transmissivity | Percentage of incoming radiation that is transmitted through the atmosphere | 0.3566 |

Supplementary Table 2: List of cultivar specific parameter values for six Chinese maize cultivars derived from experimental data obtained during the 2011 growing season (Chen et al., 2013, 2014).

| Parameters | Description | ZD958 | ZD2 | DY13 | YD13 | ND108 | XY335 |
|---|---|---|---|---|---|---|---|
| LeafNum | Leaf number | 22 | 22 | 22 | 22 | 22 | 21 |
| seedMass | Endosperm mass of the seed the plant grows from initially (mg) | 295 ± 1.75 | 291 ± 4.8 | 214 ± 1.35 | 232 ± 4.9 | 272 ± 4.75 | 312 ± 4.85 |
| $a_g$ | Intercept of grain N concentration to leaf N concentration (g N /g DW) | 0.00121 ± 0.00305 | 0.00323 ± 0.00357 | 0.0044 ± 0.00205 | 0.000188 ± 0.00282 | 0.0062 ± 0.00248 | -0.0128 ± 0.0044 |
| $b_g$ | Slope of grain N concentration to leaf N concentration ($g^{-1}$DW) | 0.778 ± 0.192 | 0.746 ± 0.206 | 0.655 ± 0.124 | 0.82 ± 0.171 | 0.529 ± 0.151 | 1.97 ± 0.321 |
| fNstem | Stem structural N concentration (g N/g DW) | 0.0038 ± 0.00017 | 0.0045 ± 0.00013 | 0.0049 ± 0.0005 | 0.0043 ± 0.00016 | 0.0038 ± 0.00017 | 0.0042 ± 0.00028 |
| $\lambda$ | Maximum photosynthesis under non-limiting leaf nitrogen ($\mu$mol/($m^2 \cdot s$)) | 24.7 ± 0.866 | 23.8 ± 0.863 | 24.1 ± 0.658 | 23.9 ± 0.868 | 21.2 ± 0.839 | 31.8 ± 0.979 |
| initD | Initial root Diameter (m) | 0.00154 ± 0.00013 | 0.00107 ± 0.000052 | 0.00104 ± 0.000034 | 0.00115 ± 0.000062 | 0.00118 ± 0.000069 | 0.00145 ± 0.000077 |
| RTD | Root tissue density ($g/dm^3$) | 90 ± 11.7 | 86 ± 6.57 | 120 ± 6.44 | 109 ± 14.4 | 101 ± 6.77 | 96 ± 7.57 |
| RDM | Ratio in diameter of mother and daughter root | 0.305 ± 0.032 | 0.396 ± 0.018 | 0.456 ± 0.017 | 0.388 ± 0.015 | 0.407 ± 0.026 | 0.318 ± 0.020 |

Supplementary Table 3: List of soil N related parameters values used for model validation as derived from experiments in 2011 (Chen et al., 2013)

| Parameter | Description | HN | MN | low N |
|---|---|---|---|---|
| Ninit | Initial soil nitrogen before N fertilizer application ($\mu$mol/m$^3$) | 3 | 3 | 3 |
| Nm | Soil N gradually released during the growing season by mineralization ($\mu$mol/m$^3$) | 6 | 6 | 6 |
| total Napp | Total N fertilizer applicated for whole growing season ($\mu$mol/m$^3$) | 5.714 | 2.857 | 0 |

Supplementary Table 4: AIC values and required number of parameters for models to relate rate of maize photosynthesis to radiation density from available data (Chen et al., 2013) allowing for more or less cultivar specific parameters. In bold the finally selected model based on the lowest AIC-value

| Model | $\alpha$ | $A_{max}$ | Rd | Number of parameters | AIC |
|---|---|---|---|---|---|
| Fit full | D* | D | D | 18 | 169.57 |
| Fit1 | S | D | D | 13 | 164.38 |
| Fit2 | D | D | S | 13 | 160.08 |
| Fit3 | D | S | D | 13 | 246.48 |
| **Fit4** | **S** | **D** | **S** | **8** | **156.41** |
| Fit5 | D | S | S | 8 | 257.40 |
| Fit6 | S | S | D | 8 | 242.95 |
| Fit7 | S | S | S | 3 | 311.17 |

* D represents different values for each cultivar while S represents same value for all cultivars
